# Predicting photosynthetic pathway from anatomy using machine learning

**DOI:** 10.1101/2023.09.11.557216

**Authors:** Ian S. Gilman, Karolina Heyduk, Carlos A. Maya-Lastra, Lillian P. Hancock, Erika J. Edwards

## Abstract

- Plants with Crassulacean acid metabolism (CAM) have long been associated with a specialized anatomy, including succulence and thick photosynthetic tissues. Firm, quantitative boundaries between non-CAM and CAM plants have yet to be established – if they indeed exist.
- Using novel computer vision software to measure anatomy, we combined new measurements with published data across flowering plants. We then used machine learning and phylogenetic comparative methods to investigate relationships between CAM and anatomy.
- We found significant differences in photosynthetic tissue anatomy between plants with differing CAM phenotypes. Machine learning based classification was over 95% accurate in differentiating CAM from non-CAM anatomy, and had over 70% recall of distinct CAM phenotypes. Phylogenetic least squares regression and threshold analyses revealed that CAM evolution was significantly correlated with increased mesophyll cell size, thicker leaves, and decreased intercellular airspace.
- Our findings suggest that machine learning may be used to aid the discovery of new CAM species and that the evolutionary trajectory from non-CAM to strong, obligate CAM requires continual anatomical specialization.

## INTRODUCTION

Carbon concentrating mechanisms increase the efficiency of photosynthesis by raising the concentration of CO_2_ inside photosynthetic tissues relative to the ambient environment. The most common carbon concentrating mechanism, Crassulacean acid metabolism (CAM), was first discovered because of marked physiological differences between succulent and nonsucculent plants (de Saussure, 1804). Generally, CAM species conduct gas exchange at night to reduce transpirational water loss; the nocturnally fixed carbon is stored as malic acid overnight and released the next day behind closed stomata, thereby saturating photosynthetic tissues with CO_2_ (Osmond, 1978). Although the co-occurrence of CAM and succulent anatomy is so consistent that botanists have used it as a guide to find new CAM plants (Coutinho, 1969), quantitative relationships between anatomy and CAM remain elusive.

CAM and succulence may be correlated because they are co-selected as adaptations to water limitation. CAM species can be up to eightfold as water use efficient as C_3_ species (Winter *et al*., 2005) and the water stored in succulent plants is essential for drought avoidance (Males, 2017). Although such a correlation does not necessarily imply that derived anatomy is a prerequisite of, or is caused by, CAM evolution, there are at least two hypothesized direct functional links between CAM and succulent anatomy. First, storage of nocturnally fixed CO_2_ as malic acid in mesophyll vacuoles may require large vacuoles in photosynthetic cells and therefore larger, succulent mesophyll cells (Zambrano *et al.,* 2014; Töpfer *et al*., 2020). Second, increased succulence in mesophyll cells may lower intercellular air space (IAS) and therefore mesophyll CO_2_ conductance (*g_m_*) (Nelson *et al*., 2008; Cousins *et al*., 2020); thus, increased succulence may increase selection for CAM by lowering the efficiency of C_3_ photosynthesis (Nelson *et al*., 2008; Edwards, 2019). It is also possible that the evolution of CAM does not entail selection on succulence *per se*, but that the use of CAM reduces constraints on succulence evolution by removing *g_m_* limitations due to carbon concentration.

Quantitative studies of CAM and anatomy have generally been restricted to relatively few taxa at the extremes of the CAM phenotypic spectrum, but have generally found positive correlations between CAM and succulence. Individual studies have reported that CAM species tend to have greater leaf thickness (LT) and larger mesophyll cell area (MA), but mixed trends have been observed for IAS (Nelson *et al*., 2005 2008; Zambrano *et al*., 2014; Earles *et al*., 2018; Luján *et al*., 2022); however, a recent meta-analysis of these relationships found inconsistent trends across clades (Herrera, 2020). Recently, hybrids between species with different photosynthetic types have been used to study the relationships between CAM activity and anatomical traits. In both *Yucca* (Agavoideae, Asparagaceae) (Heyduk *et al*., 2020) and *Cymbidium* (Orchidaceae) (Yamaga-Hatakeyama *et al*., 2022), hybrids of C_3_ x CAM crosses possessed intermediate anatomical phenotypes and CAM activity. Within *Yucca* hybrid genotypes, however, the correlations between CAM activity and anatomy decreased in magnitude or disappeared entirely (Heyduk *et al*., 2020).

The mosaic of past research provides limited insight into the evolution of CAM and photosynthetic tissue anatomy because it has focused on the extremes of CAM phenotypes (i.e., non-CAM species and species that use CAM as their primary metabolism). However, there are many recognized CAM phenotypes that differ in pattern and magnitude of CAM activity (Winter, 2019). Here, we use term “CAM” to refer to all species capable of CAM, regardless of strength or pattern of expression, and “minority-CAM” and “primary-CAM” to refer to species that fix the minority and majority of CO_2_ with CAM, respectively. Primary-CAM (pCAM) is consistent with past definitions of “CAM plant” (Winter, 2019) and “strong CAM” (Edwards, 2019), while minority-CAM (mCAM) encompasses species that can facultatively use CAM or constitutively use CAM at low levels, but primarily use C_3_ or C_4_ photosynthesis for CO_2_ assimilation (mCAM = “C_3_+CAM” of Edwards (2019), but with the inclusion of C_4_+CAM species). It is generally assumed that the evolution of pCAM requires transitioning through mCAM (Hancock and Edwards, 2014; Yang *et al*., 2015; Edwards, 2019), but the relative timings of anatomical shifts during the evolution of mCAM and pCAM – and whether or not mCAM species possess a specialized anatomy – remain open questions.

Here, we combined anatomical measurements from thousands of angiosperm species from over 200 families to draw anatomical boundaries between non-CAM, mCAM, and pCAM phenotypes. Using supervised machine learning models, we were able to classify CAM phenotypes from anatomical measurements with moderate to high accuracy. Finally, in a detailed study of the Portullugo clade (Fig. 1), we reconstructed the evolution of CAM and used phylogenetic comparative methods to establish significant relationships between anatomy and CAM evolution. Our findings support the hypothesis that CAM evolution entails anatomical evolution and reveal nuances about the earliest stages of CAM evolution.

**Figure 1.**
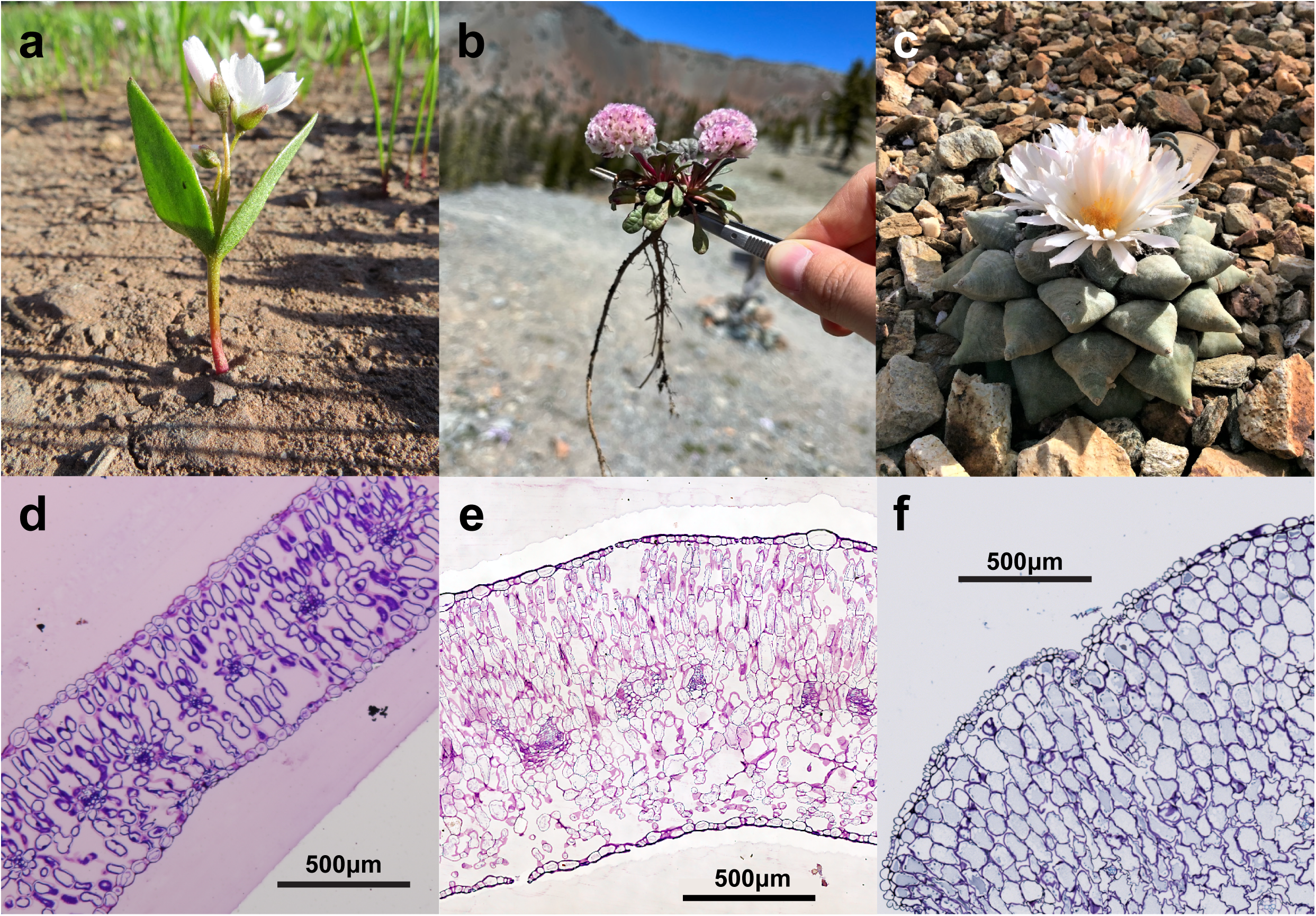
Gross morphology (a-c) and photosynthetic anatomy (d-f) of species with varying CAM phenotypes sampled for this study: (a,d) non-CAM *Claytonia lanceolata* Pursh (Montiaceae) (b,e) minority CAM *Calyptridium umbellatum* (Torr.) Greene (Montiaceae), (c,f) primary CAM *Ariocarpus retusus* Scheidw. (Cactaceae). Non-author photograph credits: (a) Dr. Thomas Stoughton, (b) Anri Chomentowska, and (c) Desert Botanic Garden, Phoenix, AZ.

## MATERIALS AND METHODS

### Public anatomical data sets, taxon sampling, and specimen imaging

Publicly available data were gathered from the TRY (Fraser *et al*., 2020) and BROT2 (Tavşanoğlu and Pausas, 2018) plant trait databases and individual studies of CAM anatomy in Orchidaceae (Silvera *et al*., 2005), Bromeliaceae (Males, 2018), Asparagaceae (Heyduk *et al*., 2016), Caryophyllales (Ogburn and Edwards, 2012, 2013), Papua New Guinean epiphytes (Earnshaw *et al*., 1987), Clusiaceae (Luján *et al*., 2022), and across angiosperms (Nelson *et al*., 2008) (Supporting Information Table S1). These data contained observations of mesophyll cell area (MA), leaf thickness (LT), mesophyll intercellular air space (IAS), leaf dry matter content (LDMC), and specific leaf area per unit dry mass (SLA). We generated two new datasets of MA, IAS, and LT for members of the Asparagaceae (subfamilies Agavoideae and Nolinoideae) and Portullugo (including families Anacampserotaceae, Cactaceae, Didiereaceae, Montiaceae, Molluginaceae, and Portulacaceae) (Supporting Information Table S2). In 2017, leaf cross sections were taken from 15 Portullugo species grown at Brown University, Providence, RI. Tissue sections were immediately placed in 10% neutral buffered formalin and sent to the Veterinary Diagnostic Laboratories in the College of Veterinary Medicine at the University of Georgia (Athens, GA) for fixation, embedding, and sectioning and staining with toluidine blue. In the spring of 2019, we collected leaf or stem cross sections of 41 species of Asparagaceae and 38 species of Portullugo growing at the Desert Botanical Garden, Phoenix, AZ; fixed specimens were created as above and imaged on an Olympus BX51 microscope (Evident Corporation, Toyko, Japan) with an Infinity3-3UR camera (Teledyne Lumenera, Ottawa, Canada). To supplement our sampling, we were provided high resolution images of leaf cross sections of 13 *Portulaca* (Portulacaceae) species used in Ocampo *et al*. (2013) by the authors.

The multiple data sets had some taxonomic overlap and some included multiple measurements from multiple accessions of the same species. To reduce our data set to one observation per species, we took the mean of each feature where multiple accessions were measured; these mean species values were used as the basis for analysis throughout. We binned each taxon into three CAM phenotypes based on Gilman *et al*. (2023) and references therein: C_3_, C_3_-C_4_, and C_4_ taxa were coded as “non-CAM”; taxa that fix the minority of their daily CO_2_ with CAM (C_3_+CAM, C_3_-C_4_+CAM, and C_4_+CAM) were coded as minority CAM (mCAM); and taxa that primarily use CAM to fix CO_2_ (i.e., over 50%, resulting in 8^13^C ratios δ-18.7‰; Winter and Holtum, 2002) as primary CAM (pCAM). The final data set contained observations from 5,316 non-CAM, 207 mCAM, and 222 pCAM taxa (Supporting Information Dataset S1).

### Measuring plant anatomy

Automated analyses of plant tissues can be difficult because many or most cells are in direct contact with other cells around much of their perimeter, rather than being separated by clear boundaries. We developed a lightweight image segmentation tool built in Python 3 with OpenCV v4.5.2 (Bradski, 2000) called MiniContourFinder to facilitate measurement of histology slides. Segmentation in MiniContourFinder is accomplished through a combination of thresholding, gradient, and morphological operations (Supporting Information Figure S1). MiniContourFinder was designed to allow users with minimal experience on the command line or image processing to quickly generate accurate and reproducible contours, particularly from plant histology images. MiniContourFinder can be run through the command line or a graphical user interface to tune contours in real time. We used MiniContourFinder to measure MA in our new Asparagaceae and Portullugo data sets. We used ImageJ v1.53 (Schneider *et al*., 2012) to calculate LT (for leafy species) and IAS (in roughly 300 μm x 300 μm areas of mesophyll).

### Statistical analysis

We investigated group differences in anatomical measurements by assessing normality and homoscedasticity, comparing raw and transformed data, testing for group differences, and finally using post-hoc tests to identify group differences. We first assessed assumptions of normality using D’Angostino and Pearson’s test (D’Agostino and Pearson, 1973) and homoscedasticity using Bartlett’s test (Bartlett, 1937) of raw and log_10_-transformed data. None of the features were normal when raw or transformed, but log_10_-transformation substantially decreased heteroscedasticity: all transformed features were homoscedastic except SLA, which was much less heteroscedastic (Supporting Information Fig. S2 and Table S3). We therefore continued with Kruskal-Wallis (KW) tests for group differences (Kruskal and Wallace, 1952) with the transformed data, and Dunn’s post-hoc tests (Dunn, 1964) where KW tests revealed significant group differences. We tested for correlations between transformed features using Pearson’s *r* (Pearson, 1895). All statistical analyses were performed using Python v3.7.12, scipy v1.5.3 (Virtanen *et al*., 2020), and scikit-posthocs v0.6.4 (Terpilowski, 2019).

### Supervised classification

We attempted to classify species’ CAM phenotypes based on anatomy using the supervised learning method gradient boosting implemented in XGBoost via the Python package ‘xgboost’ v.1.5.0 (Chen and Guestrin, 2016). XGBoost implements gradient tree boosting algorithms (Friedman *et al*., 2000; Friedman, 2001) that use greedy learning over an ensemble of regression trees to train classification models. XGBoost is rare in that it can accept observations with missing values without the need for data imputation. We conducted multiclass classification of non-CAM, mCAM, and pCAM taxa and a simpler, binary classification of non-CAM and CAM taxa, where mCAM and pCAM taxa were combined. We explored a variety of alternative parameterizations: changing the default booster (gbtree) to DART (Rashmi and Gilad-Bachrach, 2015), which can reduce overfitting by randomly dropping decision trees; changing the objective function (softmax or softprob for multiclass classification; logistic probability, logistic raw score, or hinge loss for binary classification); and changing the evaluation metric (multiclass logloss, AUC, or multiclass error rate for multiclass classification; error rate for binary classification) (the AUC evaluation metric required a softprob objective function). In all cases we randomly divided our data set between training (80%) and testing (20%).

We also tried several strategies to reduce the effects of highly imbalanced classes and sparsity. We attempted to reduce class imbalance by adjusting the parameter ‘max_delta_step’ (MDS), by random over- or under-sampling our training data, and by merging mCAM and pCAM into a binary classification model. Increasing MDS above its default (0) creates an additional penalty that reduces splitting within trees, or the addition of trees entirely, in highly imbalanced data sets. Random over-sampling (ROS) resamples minority classes until all class labels are equal (augmenting training data), while random under-sampling (RUS) subsamples classes until all class labels are equal (reducing training data). Our data were also quite sparse (67% missing data) because we merged data from largely non-overlapping studies. We evaluated three data imputation strategies: median (missing features were imputed with the median), iterative (missing features were imputed by regression of present features), and *K*-nearest neighbors (Knn; missing features were imputed using the nearest neighbors in a Knn embedding).

### Phylogenetic tree inference

The Portullugo, the clade inclusive of the Portulacineae (families Anacampserotaceae, Basellaceae, Cactaceae, Didiereaceae, Montiaceae, Portulacaceae, and Talinaceae) and its sister clade (Molluginaceae) is well-suited for large, comparative phylogenetic studies because of recent sequence data, its diversity of CAM phenotypes, and the overlap between anatomical data and extant phylogenies. We constructed a new phylogeny of the Portullugo by merging two previously published sequence matrices that were obtained using different techniques. The first dataset consisted of 841 loci from transcriptomic data used to study the evolution of Portulacineae and its adaptation to harsh environments (Wang *et al*., 2019). The second dataset was a targeted enrichment of 83 gene families, primarily with roles in plant respiration and photosynthesis (Goolsby *et al*., 2018; Hancock *et al*., 2018; Moore *et al*., 2018). To find common loci between the datasets, we independently called consensus sequences for each locus and mapped them against the sugar beet genome (assembly version EL10_1.0; McGrath *et al*., 2022) using Blast v.2.13.0 (Camacho *et al*., 2009). Mapping consensus sequences for each locus proved more accurate than using random representative sequences for a given locus due to high sequence variation. If consensus loci hit multiple reference scaffolds, we retained the reference locus with the highest bitscore. We used the resulting mapping coordinates to search for potential overlapping loci between datasets and aligned them using MAFFT v.7.508 (Katoh and Sandley, 2013). Loci showing considerable dataset overlap were concatenated to create an initial matrix of loci represented by both datasets, and then flanked with 7 randomly selected non-overlapping loci from each dataset to increase the number of taxa included and overall matrix size.

We concatenated all loci and constructed a maximum likelihood-based tree using IQ-TREE v2.2.0.3 (Minh *et al*., 2020). Within IQ-TREE, a model of sequence evolution was selected using the automated model finder (Kalyaanamoorthy *et al*., 2017) constrained to the GTR family of models; node support was assessed using ultrafast bootstrap approximation (Hoang *et al*., 2017), and the tree space was constrained by a guide tree in which all families were monophyletic. We time calibrated the tree using the fast least squares dating method (To *et al*., 2016) included in IQ-TREE using the entire concatenated sequence matrix the 13 secondary node calibrations used by Wang *et al*. (2019) from Arakaki *et al*. (2011) (Supporting Information Table S4). Confidence intervals were inferred by 100 resamplings of branch lengths by drawing new clock rates (log-normal distribution with mean 1 and standard deviation 0.2), tip dates were set to 0, and a GTR+F substitution model was selected with the automated model finder.

### Phylogenetic trait analyses

We reconstructed the evolutionary history of CAM in the Portullugo using stochastic character mapping (Nielsen, 2002; Huelsenbeck *et al*., 2003) implemented with the ‘make.simmap’ function of the R package ‘phytools’ v1.2-0 (Revell, 2012), to model CAM evolutionary history assuming 1) an all rates different (ARD) model, and 2) a constrained ARD model without reversions from pCAM to mCAM; both models assumed a root state of non-CAM. The constraint of the latter model was informed by the lack of evidence for reversions from pCAM throughout vascular plants. In all analyses, we pruned our tree to one sample per species and node reconstructions were visualized as pie charts summarizing the state frequencies over 10000 stochastic maps.

To assess the relationships between CAM phenotypes and anatomical traits in the Portullugo, we used a threshold model of trait evolution (Wright, 1934; Felsenstein, 2005), implemented with the ‘threshBayes’ function (Revell, 2014) of ‘phytools’ v1.2-0, and phylogenetic least squares (PGLS) regression (Grafen, 1989), implemented with the R package ‘nlme’ v3.1-162 (Pinheiro *et al*., 2023). We used PGLS regression to assess relationships between continuous anatomical traits and between anatomical traits and discrete CAM phenotypes (as a predictor variable). We used threshold models to measure the correlations between anatomical traits and CAM phenotype. In all analyses, our tree was pruned to match the taxa with anatomical data and reduced to one sample per taxon where necessary.

## RESULTS

Non-phylogenetic analyses of anatomy across angiosperms demonstrated significant group differences for all five anatomical features investigated (Supporting Information Table S5). Dunn’s post-hoc tests identified significant (*p* < 0.05), and generally consistent, differences between CAM phenotypes for most features: the largest differences were observed between non-CAM and pCAM phenotypes, with mCAM intermediate but not always significantly different from both non-CAM and pCAM (Fig. 2). Where sufficient data were available, these trends were supported within individual families (Supporting Information Fig. S3). We found significant negative correlations between MA and leaf dry matter content (LDMC), between LT and specific leaf area (SLA), LDMC, and IAS, and between LDMC and SLA; a significant positive correlation was found between LT and MA (Supporting Information Fig. S4).

**Figure 2.**
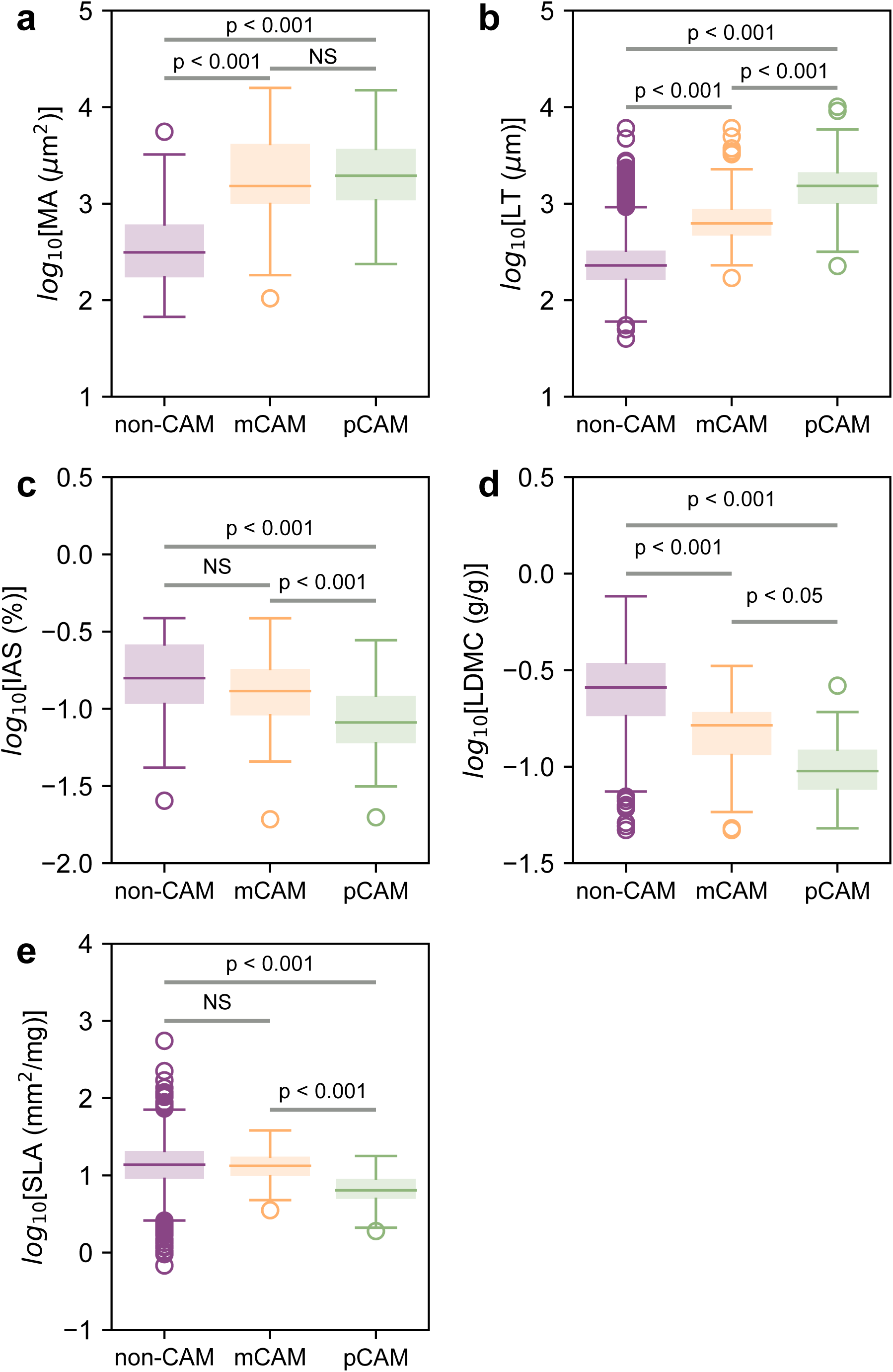
Results of Dunn’s post-hoc tests for group differences been log_10_-transformed features. Purple, yellow, and green box-and-whisker plots show non-CAM, minority CAM, and primary CAM trait distributions; boxes represent the interquartile range (IQR) with a line representing the median, whiskers show 1.5x the IQR, and points outside were considered outliers. MA, mesophyll cell area; LT, leaf thickness; IAS, intercellular airspace; LDMC, leaf dry matter content; SLA, specific leaf area.

Multiclass classification with XGBoost yielded similar results regardless of evaluation metric or objective function, with booster choice being the only source of variation (Supporting Information Fig S5). Because of the similarity of those results, we only continued using models with the softprob objective function and multiclass error rate (merror) evaluation metric (hereafter, DART and gbtree ‘base models’). The two base models had similar cross-validation test accuracies (96.0±1.1%) (Fig. 3a), precision and recall of non-CAM, mCAM, and pCAM (Fig. 3b), and feature importance rankings (LT > MA IAS ≥ LDMC > SLA) (Supporting Information Fig. S6 and Table S6). No imbalance-reduction sampling, imputation method, or alternative parameterization increased overall accuracy (Fig. 3a); however, random over-sampling (ROS) and random under-sampling (RUS) increased recall for mCAM and pCAM taxa (Fig. 3b). Between models of similar accuracy, we prioritized improving mCAM recall (also known as sensitivity in binary classification; true positives / true positives + false negatives) because true negative rates of mCAM are not well known in most CAM-evolving clades. While decreased non-CAM classification accuracy slightly decreased overall model accuracy, ROS raised recall rates of mCAM and pCAM classification to 70% and >75%, respectively. Although RUS further increased mCAM and pCAM recall (Fig. 3b), the substantial difference between training and testing accuracy (Fig. 3a) suggested that these models were overfit. To further address class imbalance, we combined mCAM and pCAM into a single “CAM” category and attempted binary classification. Binary classification models had similar test accuracies (Fig. 3c), but the hinge objective function yielded slightly higher CAM precision and recall. As in multiclass classification, ROS greatly increased CAM recall, but the F1-score (2 x precision x recall / precision + recall) remained unchanged because of an equal magnitude drop in precision (Fig. 3d).

**Figure 3.**
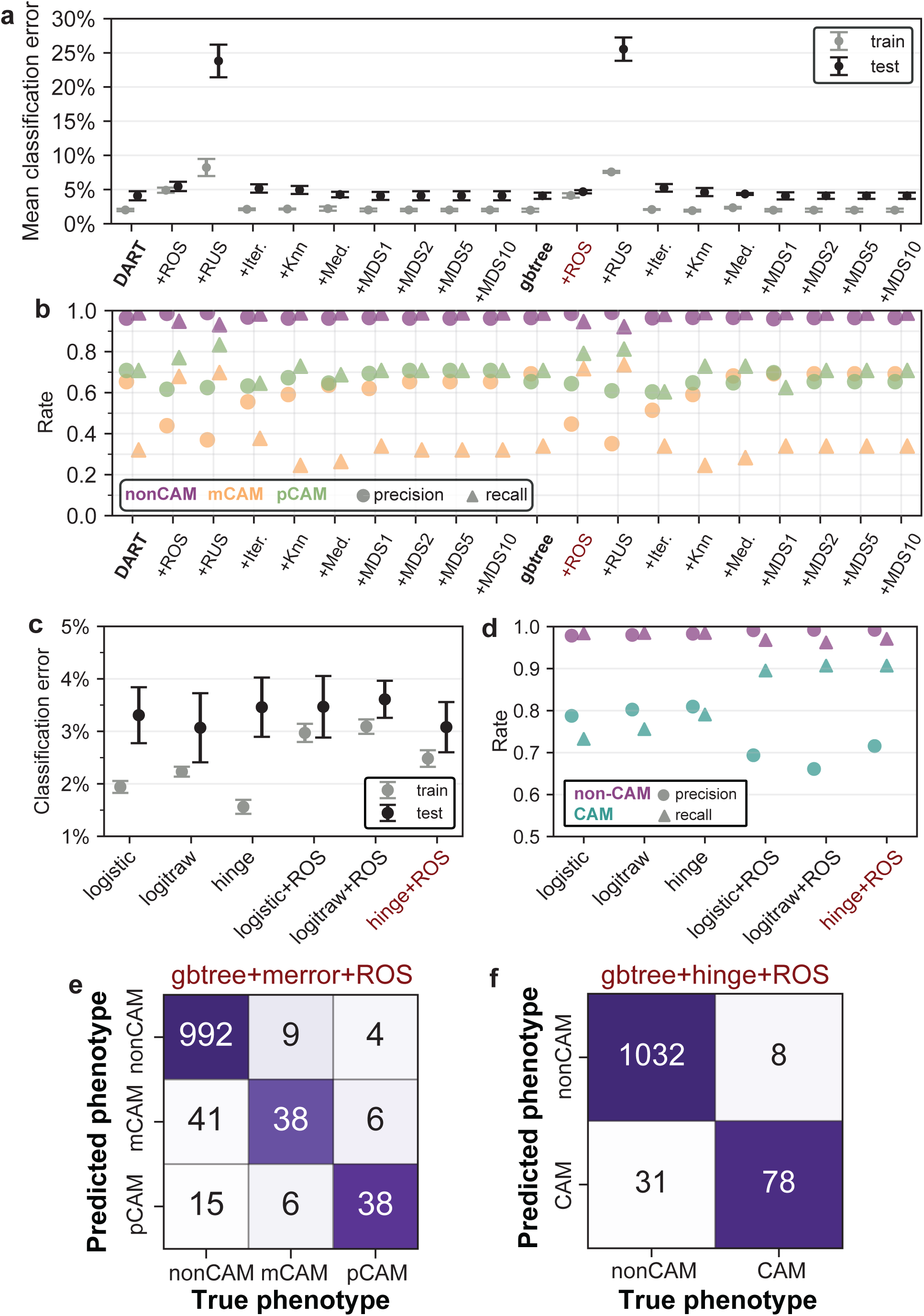
Machine learning model accuracies. Classification error (a,c), precision and recall rates (b,d), and best performing model confusion matrices (e-f) for multiclass and binary classifiers. Multiclass models (a-b) varied in booster (DART or gbtree), sampling strategy (ROS or RUS), imputation method (iterative, Knn, or median), and MDS (1, 2, 5, or 10); binary models (c-d) varied in objective function (logistic, logitraw, or hinge) and sampling strategy (with or without ROS). The columns of each confusion matrix (e-f) show the number of true CAM phenotypes in the test data set and the rows show the model predictions. The diagonal in each matrix represents correct model predictions and off-diagonal elements show incorrect predictions; for example, a true pCAM species predicted to be non-CAM would be shown in the first row, third column of (c). Knn, *K*-nearest neighbors; MDS, max_delta_step; mCAM, minority CAM; pCAM, primary CAM. Base models are in bolded text and the best performing models are highlighted in red.

Our preferred multiclass and binary classifiers both used gbtree boosters and ROS, and the hinge object function for binary classification (Fig. 3e-f). Mean cross-validation accuracies were 95.7±0.7% and 96.1±0.6% for multiclass and binary models, respectively (Fig. 3a,c). Most non-CAM taxa incorrectly classified by multiclass models belonged to clades with diverse CAM phenotypes (e.g., Bromeliaceae and Orchidaceae subfamily Epidendroideae), and mCAM taxa were roughly equally classified as non-CAM or pCAM (Fig. 3e; Supporting Information Table S7). Similarly, most incorrect predictions by the binary model were non-CAM species from CAM-evolving lineages classified as CAM (Fig. 3e; Supporting Information Table S8); generally, these taxa have not been thoroughly assessed for mCAM, and so it is possible that they may actually have a facultative or very weak CAM cycle.

Our time calibrated species tree was congruent with those from which data were compiled. Support was generally high, although multiple nodes along the backbone were unresolved and left as polytomies in downstream analyses (Fig. 4; Supporting Information Fig. S7). Stochastic character map reconstructions of CAM evolution suggested that mCAM evolved at the base of Portulacineae, and that multiple transitions to pCAM occurred in the Cactaceae and Didiereaceae, while multiple reversions to non-CAM occurred in the Montiaceae (Fig. 4).

**Figure 4.**
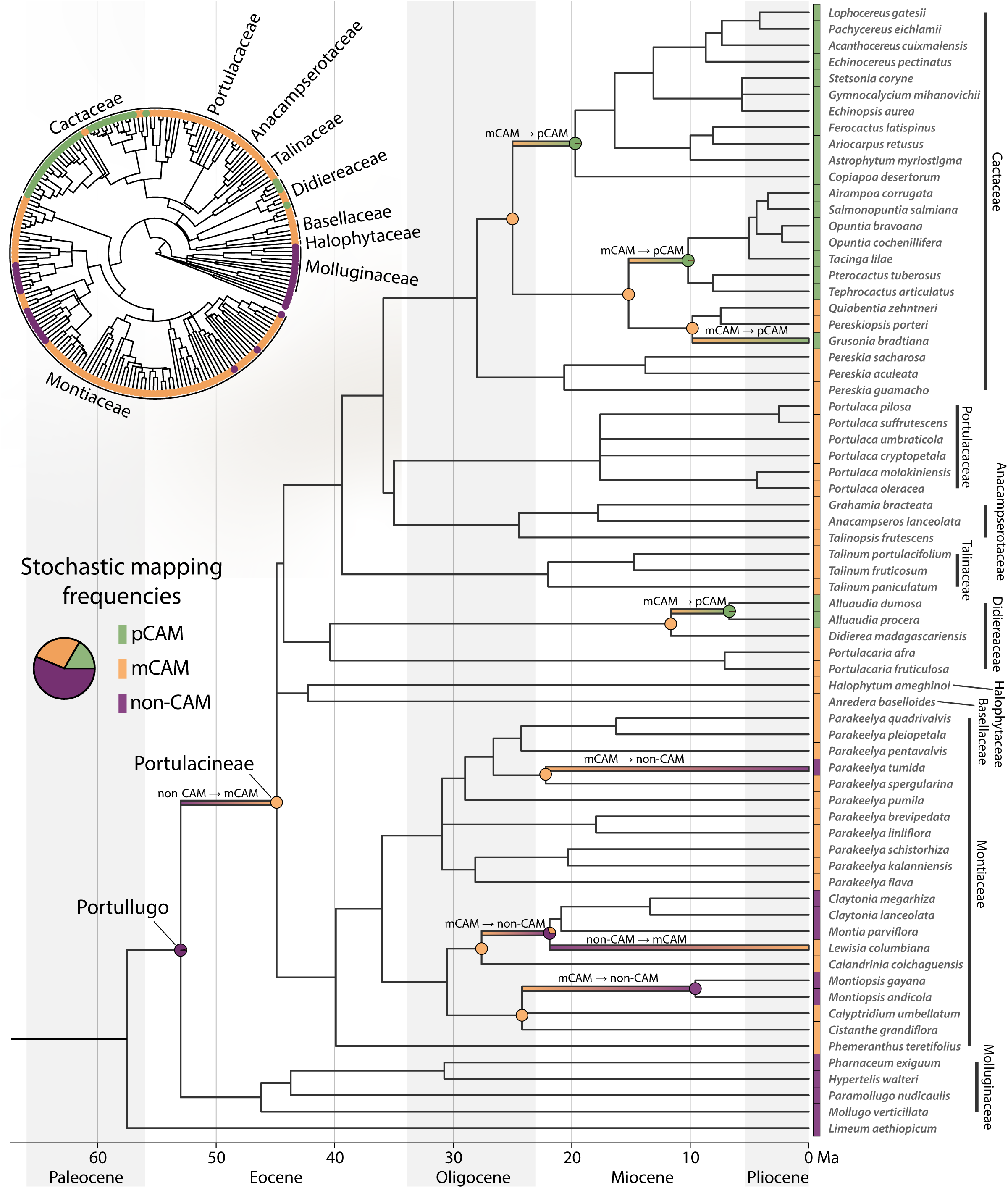
Time calibrated phylogeny of the Portullugo with inferred transitions between CAM phenotypes. The Portullugo and Portulacineae nodes are highlighted, and color gradients indicate transitions between non-CAM (purple), mCAM (yellow), and pCAM (green) based on the results of our biologically-informed ancestral state reconstruction. Pie charts at nodes bracketing inferred transitions show the fractions of stochastic maps supporting each ancestral state. This tree has been pruned to show only those taxa with morphological data used in this study and therefore not all transitions are shown; the full tree is shown in the inset and multiple ancestral state reconstructions are available in the Supporting Information.

Though similar, we preferred a constrained all rates different model of CAM evolution (Fig. 4; Supporting Information Fig. 8) to an unconstrained model (Supporting Information Fig. S9) because there is no strong empirical evidence of reversions from pCAM in any vascular plant lineage. Significant phylogenetic signal was detected in all three traits measured across the Portullugo (Supporting Information Table S9). Phylogenetic least squares (PGLS) regression revealed multiple significant (*p* < 0.05) relationships among anatomical traits and between anatomical traits and CAM phenotype (Fig. 5, Supporting Information Table S10). However, AIC-based model selection favored a model between MA and IAS with a non-significant slope, contrary to our expectation that greater mesophyll cell size would lead to lower IAS (Fig. 5a). Greater MA was a significant (*p* < 0.0001) predictor of greater LT (Fig. 5b), and we found no relationship between IAS and LT (Fig. 5c). CAM phenotype was a significant predictor of MA, LT, and IAS (Fig. 5d-f). We next used phylogenetic threshold analyses to estimate the correlations between CAM phenotype and anatomical traits under the hypothesis that there may be anatomical boundaries between CAM phenotypes. Threshold analyses mostly supported PGLS results, and recovered significant positive correlations between CAM phenotype and both MA and LT (Fig. 6a-b). However, the posterior distribution of correlation coefficients between CAM phenotype and IAS narrowly included 0 (Fig. 6c).

**Figure 5.**
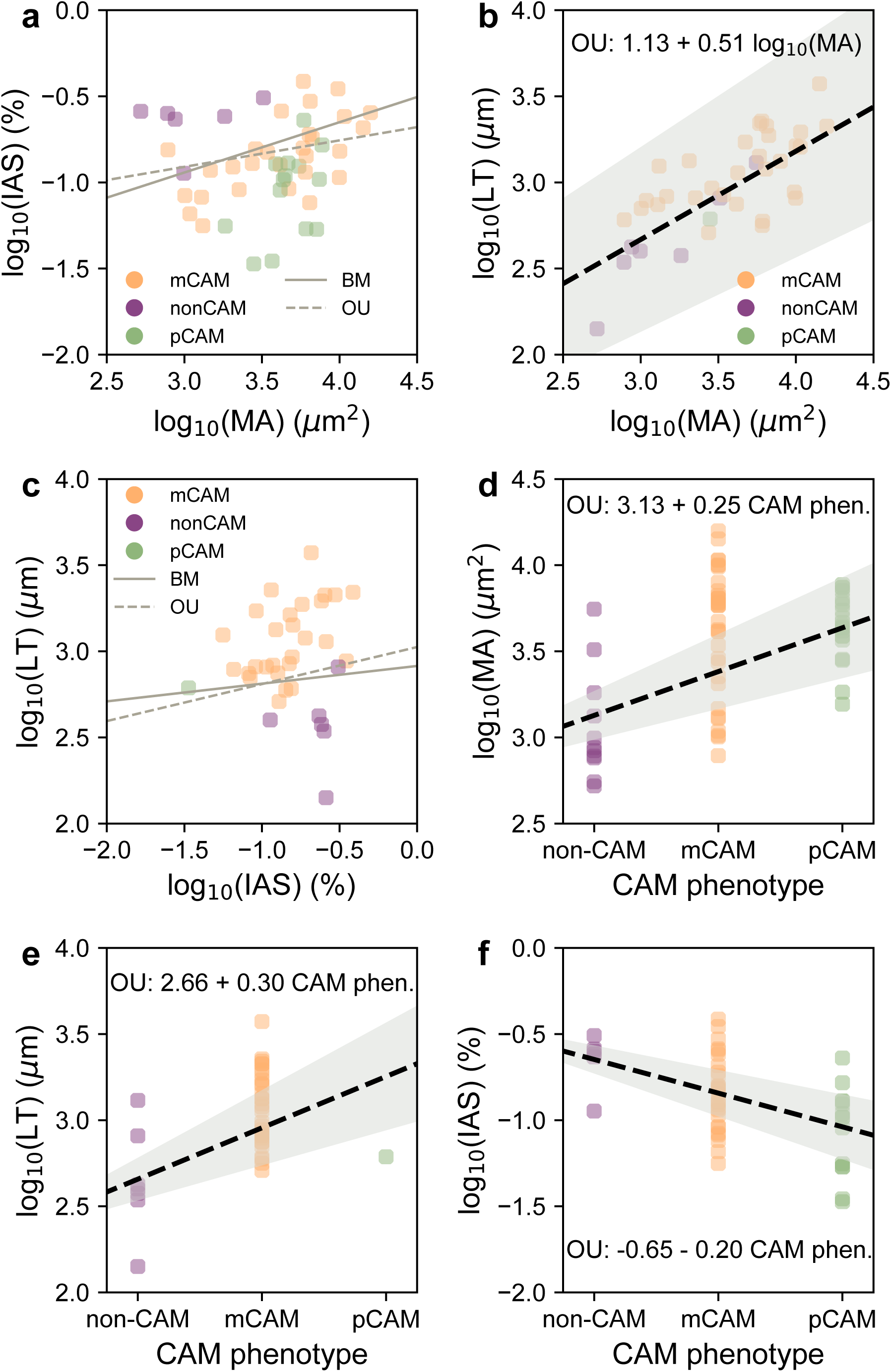
Results of phylogenetic least squares (PGLS) regression. Predictor and response variables are shown on the horizontal and vertical axes, respectively. Points show trait values for non-CAM (purple), mCAM (yellow), and pCAM (green) species. Solid and dashed grey lines show the fitted regression lines using Brownian motion (BM) and Ornstein-Uhlenbeck (OU) models of trait evolution, respectively. The best fit significant relationships are shown with bold black lines, associated model coefficients, and grey shading to show standard error. MA, mesophyll cell area; LT, leaf thickness; IAS, intercellular airspace.

**Figure 6.**
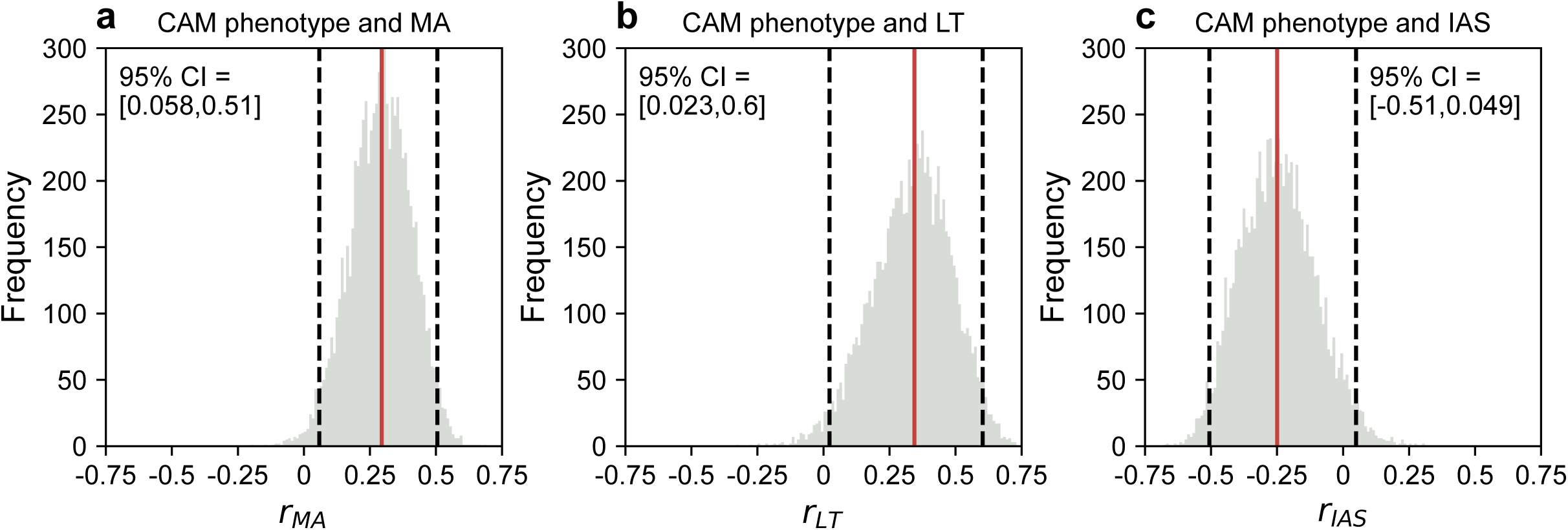
Phylogenetic threshold model correlations. The distribution of correlation coefficients (*r*) between CAM phenotype and log_10_-transformed mesophyll cell area (MA) (A), intercellular airspace (IAS) (B), and leaf thickness (LT) (C). The grey histograms show the frequency of *r* values visited by the MCMC sampler following a 20% burn-in period, red lines show the median *r* values, and dashed black lines show the 95% credible interval.

## DISCUSSION

From the beaks of Galapagos finches (Darwin, 1839) to unique inflorescence architectures (Waal *et al*., 2012), the links between form and function have always inspired biologists. Fixed in place, with passive mechanisms for carbon and water acquisition, plants rely on anatomical innovations to adapt to different environments. Succulence has long been understood as a drought avoidance adaptation, but its relationship with CAM has not been resolved as causal or merely coincidental. Through our broad survey of angiosperms and detailed study of the Portullugo, we found support for previous hypotheses of CAM and photosynthetic tissue anatomy co-evolution. Furthermore, we demonstrate that the presence or absence of CAM may be predicted using only a handful of anatomical measurements.

Anatomical measurements from over 200 angiosperm families revealed significant differences in photosynthetic tissue anatomy of non-CAM, mCAM, and pCAM species. The larger mesophyll cell area (MA) of both mCAM and pCAM species suggests some anatomical specialization is required to perform CAM in any capacity, and the reduction in intercellular airspace (IAS) of pCAM species indicates that further specialization is required to use CAM for primary carbon metabolism. We also showed significant increases in leaf thickness (LT) and decreases in leaf dry matter content (LDMC) from non-CAM to mCAM to pCAM, as well as significantly lower specific leaf area (SLA) in pCAM species, which support past anatomical studies that found thicker and more succulent leaves to be positively associated with strong CAM activity within individual clades (Teeri *et al*., 1981; Winter *et al*., 1983; Nelson *et al*., 2005, 2008; Zambrano *et al*., 2014; Luján *et al*., 2022).

Because lineage-specific organismal detail will surely influence physiology-anatomy relationships, analyses of anatomy and CAM evolution are best evaluated using phylogenetic comparative methods. PGLS regression and phylogenetic threshold analysis supported the correlated evolution of larger mesophyll cells and thicker leaves. Although PGLS regression further showed a continuous decrease in IAS from non-CAM to pCAM species, we found no significant relationship between IAS and MA. That IAS and MA may evolve independently of one another provides an important nuance to the co-evolution of succulence and CAM. Decreased IAS in CAM species has often been discussed as an adaptation to reduce CO_2_ efflux during malate decarboxylation (Nelson *et al*., 2008) or as a consequence of increased succulence restricting *g_m_*, which would limit CO_2_ fixation by Rubisco during the day (Zambrano *et al*., 2014; Earles *et al*., 2018; Edwards, 2019). More recently, reduced IAS has been hypothesized to be an indirect consequence of increased mesophyll cell volume used for malic acid storage (Leverett *et al*., 2023). While we found that succulence generally increased with CAM evolution, the decoupling of the underlying traits may allow the evolution of intermediate photosynthetic and anatomical phenotypes that efficiently utilize both CAM and C_3_ or C_4_ photosynthesis. These conclusions are consistent with photosynthetic models that found increased vacuolar volume (and therefore MA) necessary for CAM (Töpfer *et al*., 2020) and empirical findings that the high IAS in mCAM species may allow for C_3_ (or C_4_) photosynthesis when plants are not engaging CAM (Nelson *et al*., 2008; Zambrano *et al*., 2014). Furthermore, lowest IAS values in the Portullugo were observed in pCAM species, which reinforces the hypothesis that extremely low IAS may reduce *g_m_* and C_3_ or C_4_ efficiency.

In addition to providing support for a positive relationship between CAM and succulence, our findings point towards interactions between life history, CAM, and succulence for those taxa that do not neatly fall along regression lines. Phylogenetic analyses of the Portullugo showed general increases in succulence and a tightening of the distributions of underlying traits for pCAM species. In contrast, mCAM taxa had both the single largest MA and greatest IAS observations, with values that mostly spanned the non-CAM to pCAM range. The eight largest observed MA values in the Portullugo were from mCAM species; most are annual species, with the exceptions of *Parakeelya flava* (a perennial geophyte with above ground tissue that regrows annually) and *Grahamia bracteata* and *Talinopsis frutescens* (which have non-succulent woody stems and drought-deciduous leaves). This suggests that the evolution of pCAM requires a shift to a (functional) perennial life history with long-lived photosynthetic tissues (Hancock *et al*., 2019); indeed, we are unaware of any annual pCAM species. The halophyte *Halophytum ameghinoi* had the second largest observed MA in the Portullugo. While saline soils may select for increased succulence to maintain cytosolic ion balance (Naidoo and Rughunanan, 1990; Ogburn and Edwards, 2010), high salt concentrations inhibit the central CAM enzymes phosphoenolpyruvate carboxylase (PEPC) and malic enzyme (ME) (Kluge and Ting, 1978), and may therefore represent an ecological constraint on the evolution of pCAM.

Our ancestral state reconstruction of CAM in the Portullugo was the first to model CAM as an ordered multistate trait, and supported an early-to mid-Eocene origin of mCAM – a time when the Earth’s atmosphere had relatively high levels of CO_2_ (Rae *et al*., 2021). The reconstruction of mCAM at the crown of the Portulacineae agrees with transcriptomic data that suggest a single recruitment event of a PEPC ortholog for use in CAM (Christin *et al*., 2014; Goolsby *et al*., 2018). All transitions to pCAM were found be within the past 30 Ma (Sage *et al*., 2023), congruent with shifts across angiosperms (including within Caryophyllales) to C_4_ photosynthesis, as atmospheric CO_2_ fell below 500ppm (Christin *et al*., 2011). Despite declining CO_2_ in the Oligocene and Miocene, multiple lineages within the Montiaceae have lost the ability perform CAM. Although we expect more Montiaceae lineages to exhibit CAM upon experimentation, multiple independent losses of CAM have been experimentally validated in *Parakeelya* (Hancock *et al*., 2019), a clade endemic to hot, dry areas of Australia. While life history may constrain the evolution of pCAM, it remains unclear why some members of the Portulacineae – occupying similarly semiarid environments – transitioned to C_3_ photosynthesis, while others simultaneously transitioned to pCAM, as CO_2_ continued to decline. We suspect that these losses of CAM may be linked to shifts in phenology; for example, C_3_ *Parakeelya* tend to germinate toward the end of the wet season, when temperature are cooler and water is readily available.

Most clades with CAM lineages show highly bimodal distributions of carbon isotope ratios (Messerschmid *et al*., 2021) that have been used for decades to identify pCAM species, but are generally unable to distinguish between mCAM from non-CAM. Laborious controlled experiments (e.g., of gas exchange or malic acid content) with live plants have been the only ways to identify mCAM, but such experiments are not feasible for many long-lived, rare, or difficult to cultivate species. We found that differences in photosynthetic anatomy across angiosperms translated into moderate to high accuracy in predicting CAM phenotype. After assessing a variety of machine learning models, we found that random-over sampling (ROS) increased prediction of mCAM and pCAM species while not overfitting to training data. To our knowledge, machine learning has not yet been applied to predict the presence or absence of physiological traits from anatomical measurements, such as CAM phenotypes. We believe that the accuracy we obtained represents a lower bound on the true accuracy of our models because some of our misclassified non-CAM species have not been thoroughly investigated for mCAM. For example, multiple orchid and bromeliad species labeled as non-CAM were predicted to be mCAM, but have not been subjected to drought experiments that might induce CAM activity. We predict that some misclassified species, such as *Nolina bigelovii* (Asparagaceae) will exhibit CAM upon experimentation, resulting in more true positive predictions. Experimentation should continue to be the gold-standard for determining CAM phenotypes, but machine learning models, such as those developed here, could play a valuable role in prioritizing study species and would only require small tissue sections for initial fixation and measurement.

Applications of machine learning in Plant Physiologyogy and evolution are only just beginning. Machine learning has been successful in predicting real-time photosynthetic status; for example, deep learning using hyperspectral reflectance in wheat has been used to predict electron transport rate, CO_2_ assimilate rate, stomatal conductance, and more (Furbank *et al*., 2021). Our machine learning models were limited in several ways; perhaps most by the degree of missing data and class imbalance. Our greatest model improvements came when using ROS, suggesting that measuring new mCAM and pCAM species to reduce class imbalance will increase model accuracy. If missing data could be sufficiently reduced, imputation strategies may facilitate the use of models beyond XGBoost, which allows missing data. In addition to our machine learning models, we hope that the tools and methodology developed here for measuring anatomy and merging sequence matrices will facilitate future studies of anatomical evolution.

Although software exists for taking measurements from images (e.g., ImageJ (Schneider *et al*., 2012), which we used for portions of this study), making dozens or hundreds of measurements needed for phylogenetic studies remains time consuming and the results are not easily reproducible. Our image segmentation software, MiniContourFinder can be automated from the command line, quickly segment and measure image features, and record associated metadata so exact measurements can be reproduced. Finally, our strategy for combining reduced-genomic sampling data types into a single phylogenetic analysis is flexible and in theory adaptable to any sequencing strategy. Most clades have reference, or near-reference, quality genomes within ∼75 Ma of their focal taxa (as in this study) (Cheng *et al*., 2018) that can serve as common maps to identify overlapping genomic regions, and high-quality transcriptomes (Matasci *et al*., 2014; Leebens-Mack *et al*., 2019) or targeted sequencing data (Johnson *et al*., 2019) for constructing backbones in larger phylogenies.

In conclusion, with a broad sampling of anatomical traits from thousands of angiosperms and a detailed phylogenetic study of the Portullugo clade, we provided support for hypotheses of CAM anatomical evolution. Our findings suggest that even weakly expressed CAM is correlated with larger mesophyll cells, and that decreased intercellular airspace in photosynthetic tissue is associated with a transition to using CAM as the primary carbon fixation pathway. Furthermore, our findings point towards possible evolutionary constraints on pCAM evolution, such as annual life history and halophytism. We were able predict CAM phenotypes from a handful of anatomical features, which represents a successful first application of machine learning to this problem, but also highlights the paucity of anatomical data for species capable of weak or facultative CAM. As data accumulate, we hope that these correlations will be continuously evaluated across vascular plants with tools that may allow causal evolutionary inference, such as phylogenetic path analysis (von Hardenberg and Gonzalez-Voyer, 2013). We expect that efforts to quantify key anatomical parameters for a diversity of CAM phenotypes will more sharply delineate the anatomical requirements of even a weak CAM cycle, and demonstrate the anatomical and biochemical interplay during the evolutionary transition to a primary CAM physiology.

## ACKNOWLEDGMENTS

We would like to thank Dr. Jonathan Koss for advice on implementing computer vision algorithms, Dr. Elena Voznesenskaya for providing images of *Portulaca* species, Dr. Eric Lazo-Wasem and Lourdes Rojas for help imaging specimens, and Joni Ward, Rual Puente-Martinez, and the rest of the staff at the Desert Botanical Garden (Phoenix, AZ) for assistance with the living collections that contributed to this manuscript. This research was funded by the National Science Foundation (IOS-1754662 to EJE).

## AUTHOR CONTRIBUTIONS

ISG, KH, and EJE designed the research plan; ISG, KH, and LPH collected, fixed, and imaged specimens; ISG developed image segmentation software, curated morphological data, time calibrated the phylogeny, conducted statistical analyses, and wrote the first draft of the manuscript; CAM-L designed the sequence merging strategy, generated sequence matrices, and constructed the initial phylogeny; all authors contributed to editing and revising the manuscript.

## DATA AVAILABILITY STATEMENT

The data that supports the findings of this study are available in the Supporting Information of this article. All statistical analyses of this study can be found at https://github.com/isgilman/Predicting-CAM and all installation and documentation for MiniContourFinder can be found at https://minicontourfinder.readthedocs.io/en/latest/.

Additional Supporting Information may be found online in the supporting information section at the end of the article.

## SUPPORTING INFORMATION

**Figure S1** Image processing by MiniContourFinder

**Figure S2** Comparisons of raw and log_10_-transformed data.

**Figure S3** Results of Dunn’s post-hoc tests for group differences been log_10_-transformed features for select families.

**Figure S4** Correlations between log_10_-transformed features.

**Figure S5** Accuracies of base models.

**Figure S6** Relative feature importance scores.

**Figure S7** Time calibrated phylogeny of the Portullugo.

**Figure S8** Portullugo CAM constrained ARD reconstruction.

**Figure S9** Portullugo CAM ARD reconstruction.

**Table S1** Final anatomical data set information.

**Table S2** List of accessions sampled from for this study.

**Table S3** Results of D’Angostino and Pearson’s test for normality and Bartlett’s test for homoscedasticity of raw and log_10_-transformed data.

**Table S4** Node calibrations used from Arakaki *et al*. (2011).

**Table S5** Results of Kruskal–Wallis tests for group difference between CAM phenotypes.

**Table S6** Relative feature importance of multiclass models.

**Table S7** Incorrect predictions of the best performing multiclass model.

**Table S8** Incorrect predictions of the best performing binary model.

**Table S9** Phylogenetic signal in anatomical features of the Portullugo.

**Table S10** Results of phylogenetic least squares (PGLS) regressions.

**Methods S1** Summary of MiniContourFinder image segmentation algorithm.

**Dataset S1** Species’ anatomical data mean values.

