## Supplementary Information for "Predicting photosynthetic pathway from anatomy using machine learning"

**Fig. S1** Image processing by MiniContourFinder

An image of a histological section of a stem of *Alluaudia Dumosa* (Drake) Drake (a) is denoised, converted to grayscale, and undergoes adaptive histogram normalization (b). The image is adaptively blurred (c) and an adaptive Gaussian threshold is applied (d). A Laplacian operator acts as a high pass filter (e), and the image is dilated (f). The gradient is taken (g) and the result is binarized (h). Finally, the background is cleaned through morphological opening (i) and closing (j), and the image is flooded from the outside to remove partial shapes (k). Contours (magenta) are then detected and projected over the original image (l).

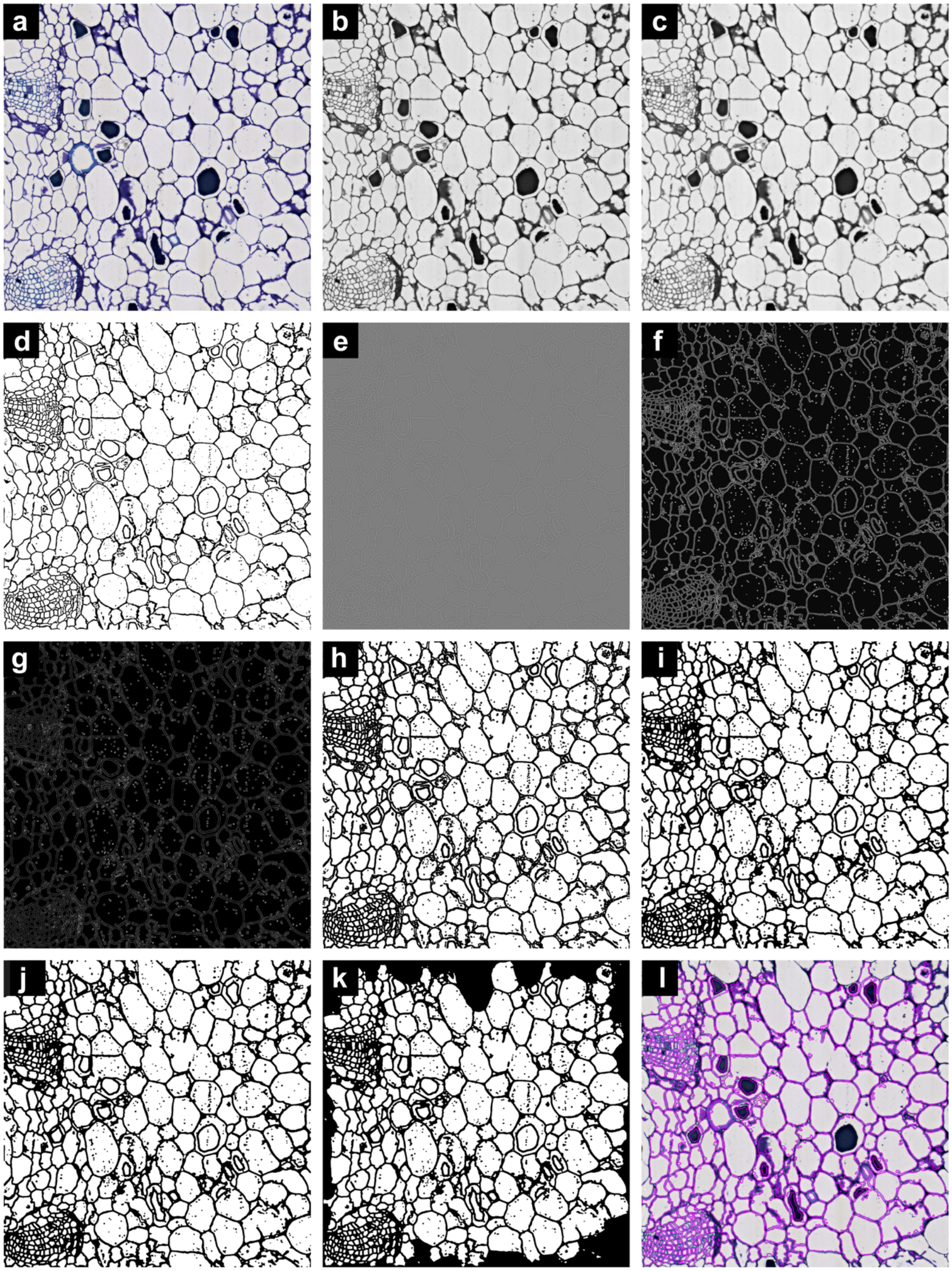


**Fig. S2** Comparisons of raw and $log_{10}$-transformed data.

Histograms of raw (a, e, i, m, and q) and $\log_{10}$-transformed data (b, f, j, n, and r), normalized frequencies of $\log_{10}$-transformed data (c, g, k, o, and s), and raw and $\log_{10}$-transformed standard deviations of each feature (d, h, l, p, t) by CAM phenotype. MA, mesophyll cell area (a-d); LT, leaf thickness (e-h); IAS, intercellular airspace (i-l); LDMC, leaf dry matter content (m-p); SLA, specific leaf area (q-t). In all plots, non-CAM, mCAM, and pCAM data are shown in purple, yellow (with hatching in histograms), and green, respectively. Raw and transformed standard deviations are represented by "X" and "O" markers, respectively (d, h, l, p, t).

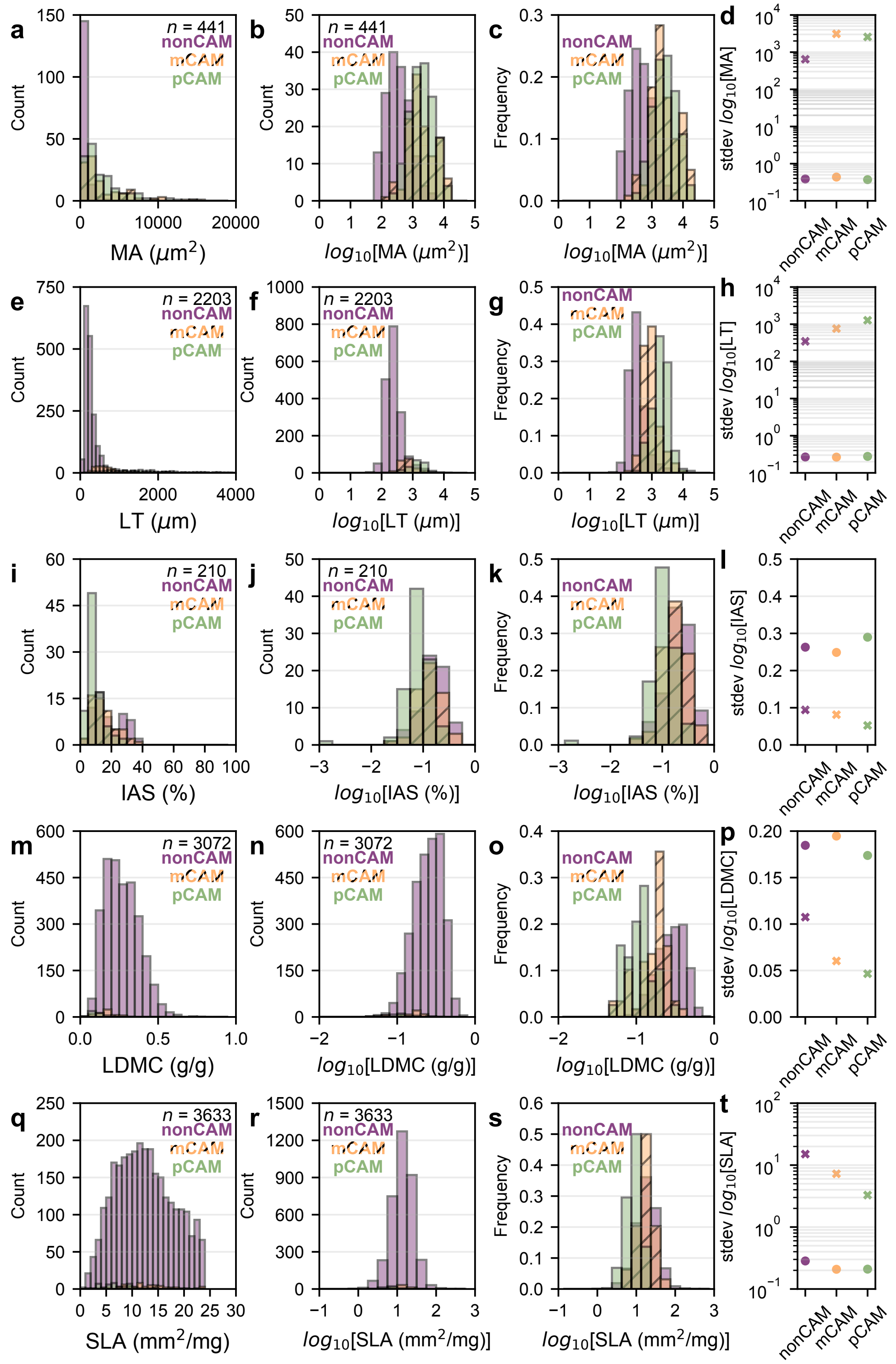


**Fig. S3** Results of Dunn's post-hoc tests for group differences been $log_{10}$-transformed features for select families.

Purple, yellow, and green box-and-whisker plots show non-CAM, minority CAM (mCAM), and primary CAM (pCAM) trait distributions; boxes represent the interquartile range (IQR) with a line representing the median, whiskers show 1.5x the IQR, and points outside were considered outliers. Tests were only conducted if there were ten or more observations of a particular CAM phenotype. MA, mesophyll cell area; LT, leaf thickness; IAS, intercellular airspace; LDMC, leaf dry matter content; SLA, specific leaf area.

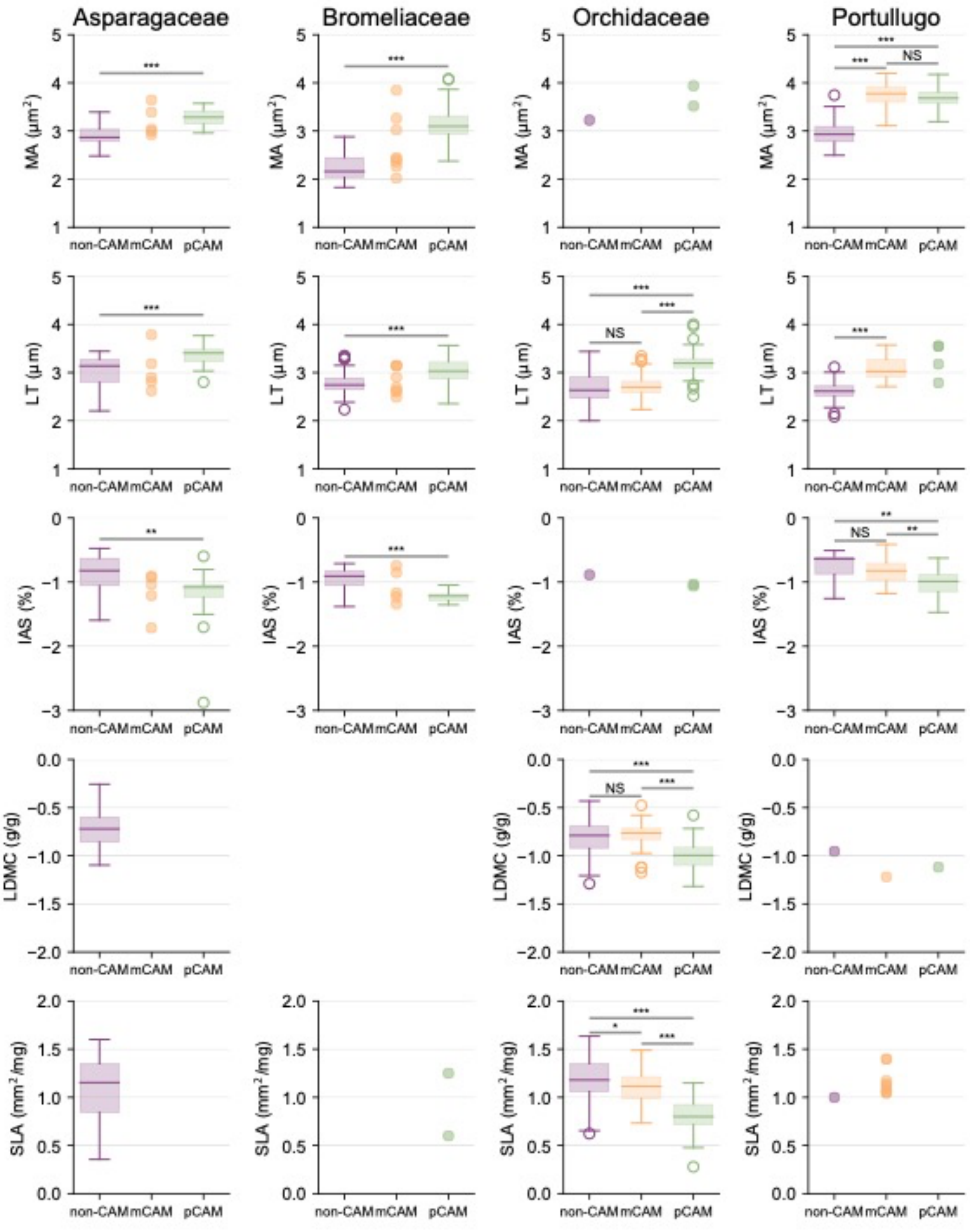


**Fig. S4** Correlations between $\log_{10}$-transformed features.

Correlations between $\log_{10}$-transformed mesophyll cell area (MA), leaf thickness (LT), intercellular airspace (IAS), leaf dry matter content (LDMC), and specific leaf area (SLA). The upper triangle of the matrix shows the correlation sign and magnitude, with strong positive correlations in red and strong negative correlations in blue, while the lower triangle indicates the significance level of the correlation; “***”, $p < 0.001$; “**”, $p < 0.01$; “*” $p < 0.05$; “NS”, non-significant

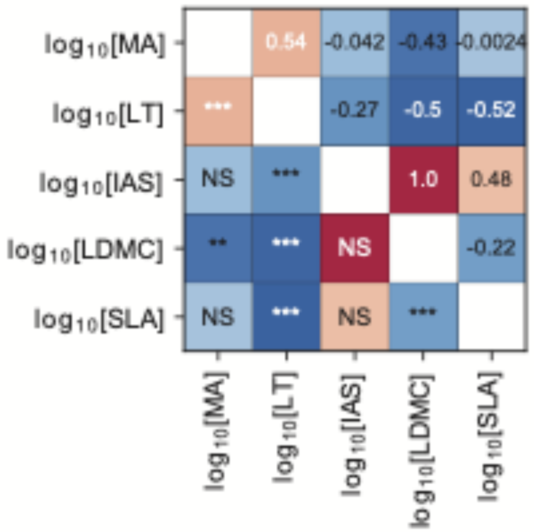


**Fig. S5** Accuracies of base models.

Confusion matrices showing the accuracies of model predictions of test data (a-j). The columns of each confusion matrix show the number of true CAM phenotypes in the test data set and the rows show the model predictions. The diagonal in each matrix represents correct model predictions and off-diagonal elements show incorrect predictions; for example, a true pCAM species predicted to be non-CAM would be shown in the first row, third column. Each of the ten models varied in booster (DART or gbtree), objective function (softmax or softprob), and evaluation metric (logloss, multiclass error, or AUC); log(L), logloss; merror, multiclass error; XGB, XGBoost; mCAM, minority CAM; pCAM, primary CAM. Model error rates are shown using merror (k), one-vs-rest area under the receiver-operator curve (AUC) (l), and multiclass logloss (m). Models using merror and logloss could use either softprob or softmax objective functions, but AUC required softprob.

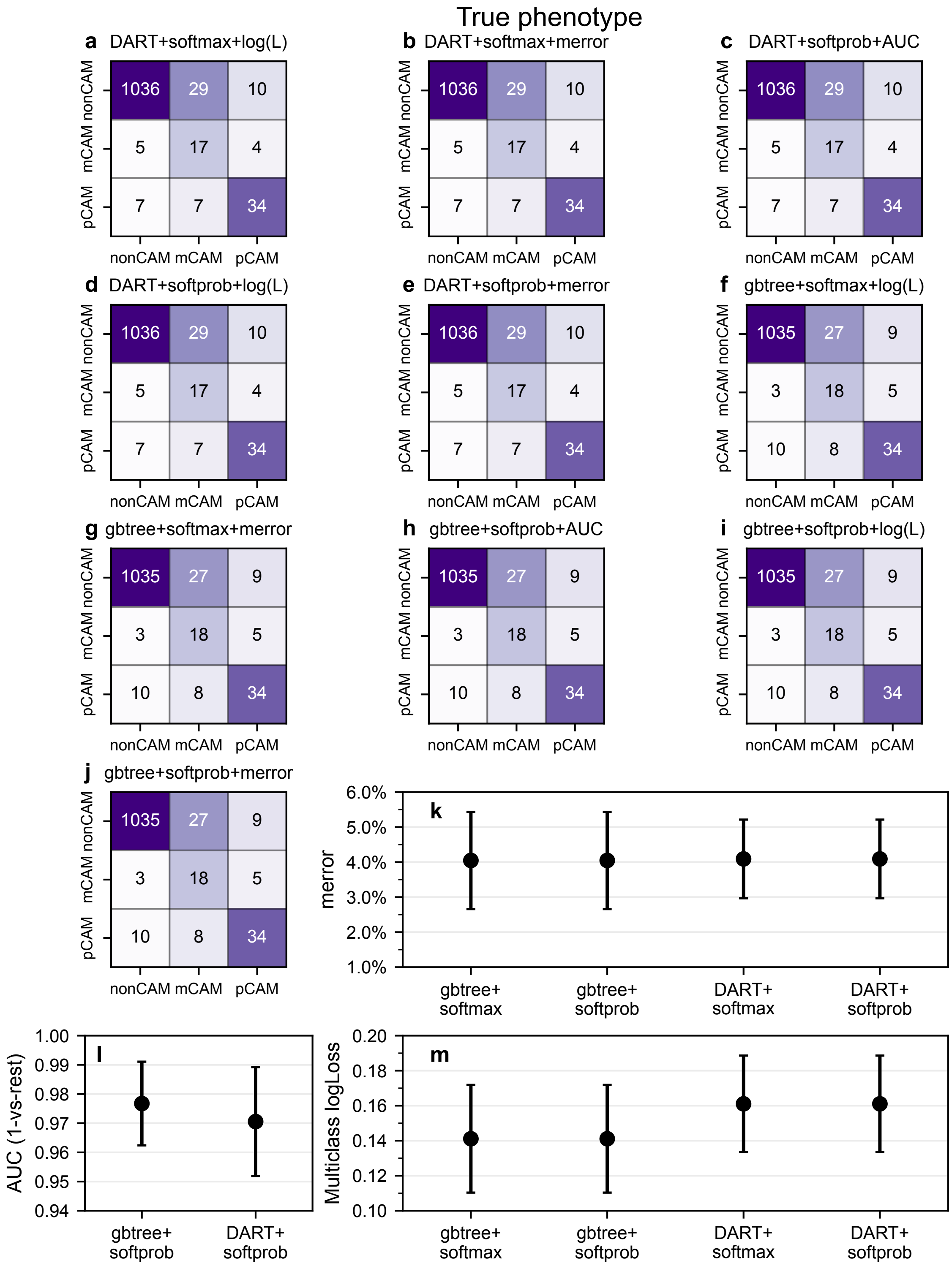


**Fig. S6** Relative feature importance scores.

Relative feature importance for models with alternative boosters (DART or gbtree), objective functions (softprob or softmax), and evaluation metrics (merror, logloss, or AUC). Base models are in bolded text and the best performing model is highlighted in red.

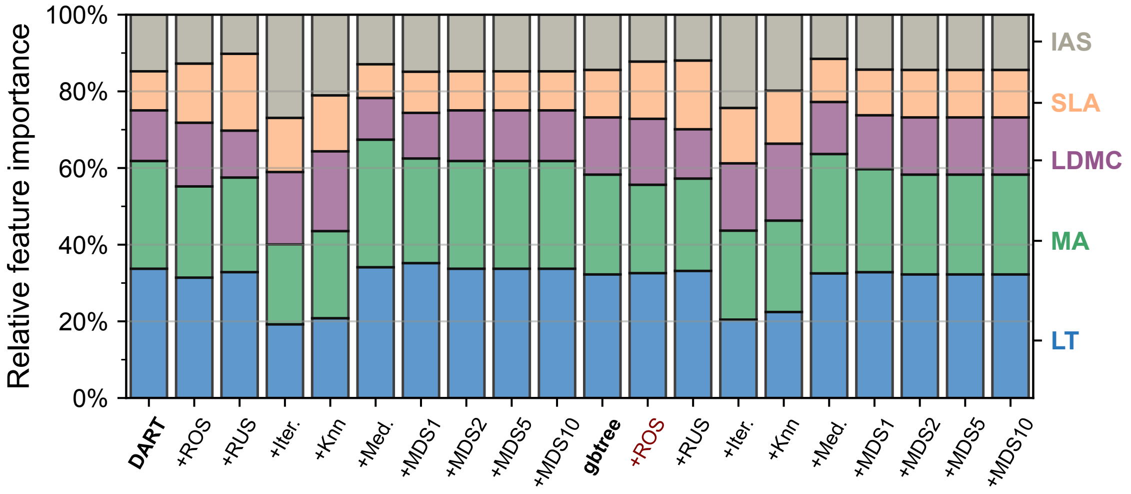


**Fig. S7** Time calibrated phylogeny of the Portullugo.

Time calibrated phylogeny of the Portullugo. Node age confidence intervals are shown in blue for bifurcating branches and calibration nodes are highlighted in yellow. Anacampserotaceae + Cactaceae + Portulacaceae (ACP); ACP + Talinaceae (ACPT)

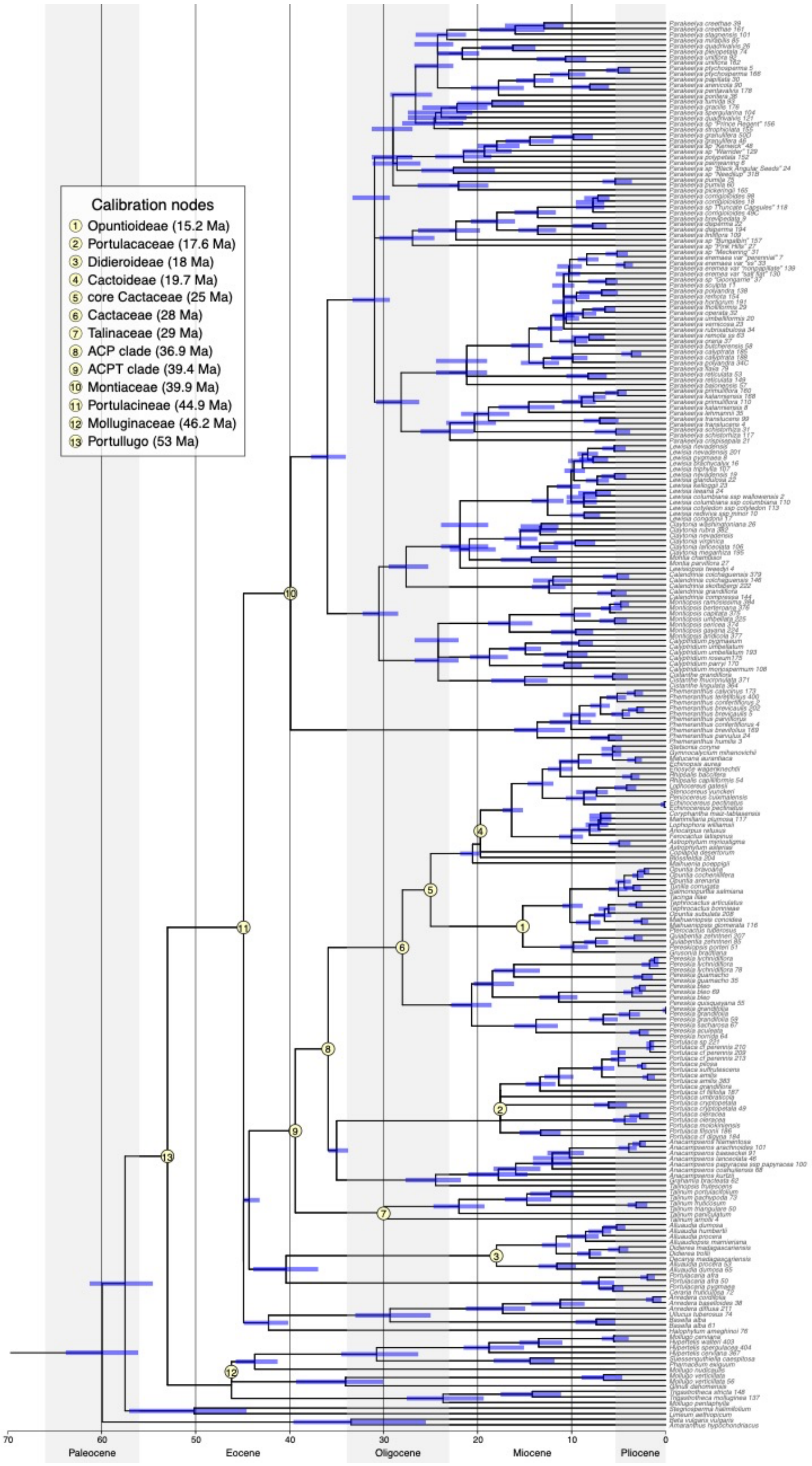


**Fig. S8** Portullugo CAM constrained ARD reconstruction.

Reconstruction of CAM evolution in the Portullugo based on 10000 stochastic character maps and an all-rates different model of evolution constrained to prevent reversions from pCAM to mCAM. Transition rates are given per million years; non-CAM, mCAM, and pCAM are shown in purple, yellow, and green, respectively; transitions are highlights with gradient-colored branches.

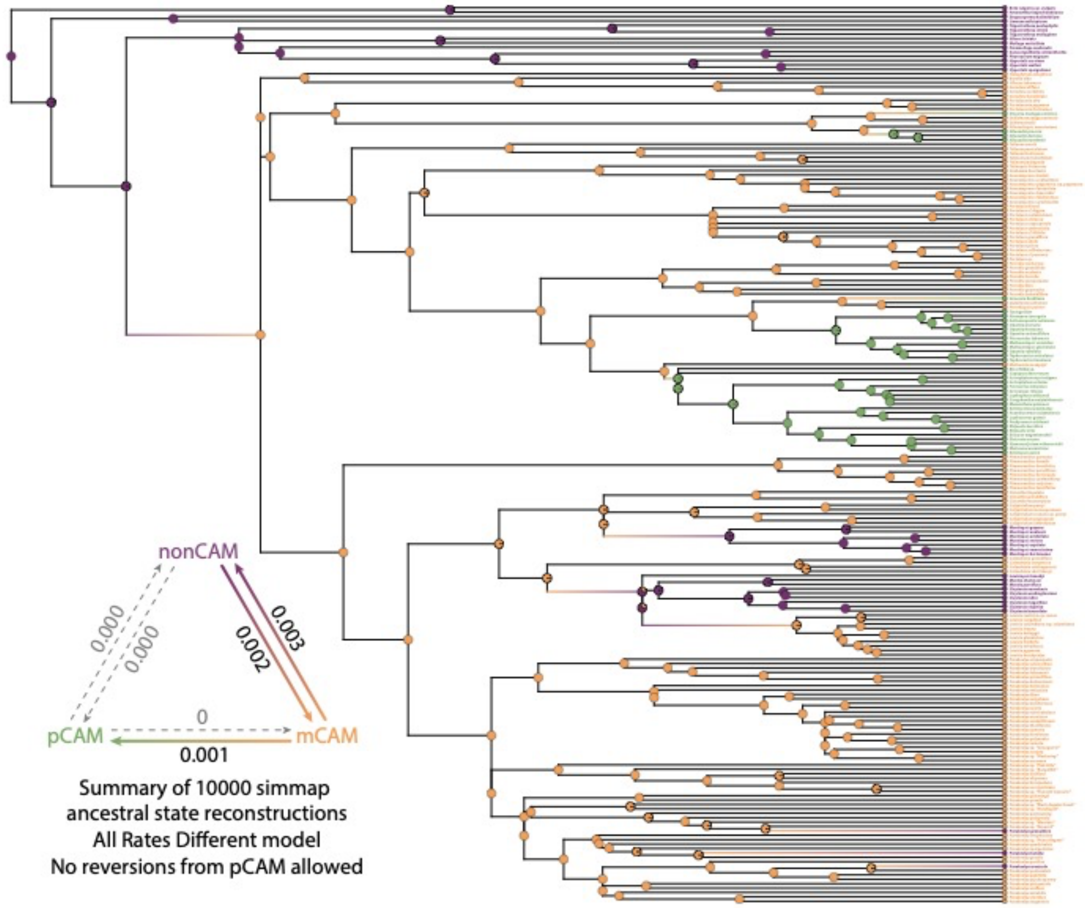


**Fig. S9** Portullugo CAM ARD reconstruction.

Reconstruction of CAM evolution in the Portullugo based on 10000 stochastic character maps and an all-rates different model of evolution. Transition rates are given per million years; non-CAM, mCAM, and pCAM are shown in purple, yellow, and green, respectively; transitions are highlights with gradient-colored branches.

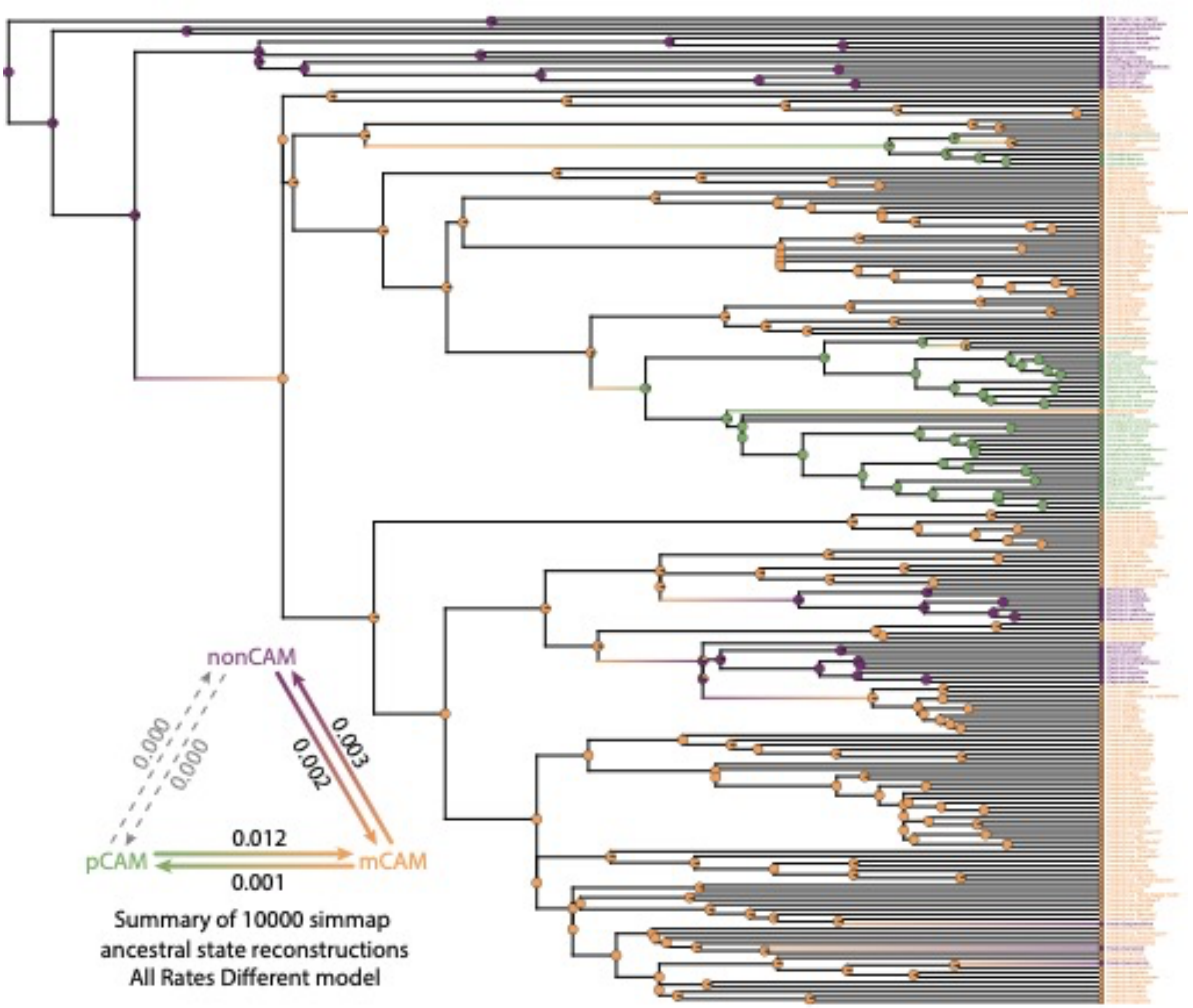


**Table S1** Final anatomical data set information..

Note that not all traits were measured for every taxon in each data set

| Study taxa | Non-CAM taxa | Minority CAM taxa | Primary CAM taxa | Traits measured |
|---|---|---|---|---|
| Asparagaceae (Heyduk *et al.*, 2016) | 4 | 2 | 7 | IAS, LT, and MA |
| Bromeliaceae (Males, 2017) | 67 | 8 | 84 | IAS, LT, and MA[a] |
| Orchidaceae (Silvera *et al.*, 2005) | 101 | 52 | 43 | SLA, LDMC[b], and LT |
| Papua New Guinea epiphytes (Earnshaw *et al.*, 1987) | 24 | 19 | 15 | LT |

| | | | | |
|---|---|---|---|---|
| Angiosperms (Fraser, 2020) | 4,718 | 21 | 8 | IAS, LDMC, LT, MA, and SLA |
| Angiosperms (Tavşanoğlu and Pausas, 2018) | 611 | 2 | 0 | LDMC and SLA |
| Angiosperms (Nelson and Sage, 2008) | 8 | 5 | 14 | IAS, LT, and MA |
| Caryophyllales[c] (Ogburn and Edwards, 2012, 2013) | 19 | 29 | 2 | IAS, LT, and MA |
| Clusiaceae (Luján *et al.*, 2022) | 9 | 55 | 1 | LT and MA |
| Asparagaceae and Portullugo[d] | 13 | 34 | 61 | IAS, LT, and MA |
| Total[e] | 5,316 | 207 | 222 | |

[a]Mesophyll cell area was calculated from chlorenchyma cell diameter assuming a circular geometry.
[b]Leaf dry matter content was calculated as the inverse of fresh mass to dry mass ratio.
[c]Some taxa were remeasured for this study.
[d]This study, including newly measured image of *Portulaca* from Ocampo *et al.* (2013), considered as mCAM here.
[e]Totals are generally lower than the sum of the studied taxa because of overlap between studies.

**Table S2** List of accessions sampled from for this study.

Samples collected from the Desert Botanical Garden (Phoenix, AZ, USA) begin with "DBG"; all remaining samples were taken from the living collections of Dr. Erika J. Edwards.

| Taxon as collected | Current taxonomy | Accession number |
|---|---|---|
| *Agave americana* L. | *Agave americana* L. | DBG 1967-9016-02-11 |
| *Agave americana* L. | *Agave americana* L. | DBG 1975-0020-01 |
| *Agave americana* L. | *Agave americana* L. | DBG 1975-0230-02-3 |
| *Agave attenuata* Salm-Dyck | *Agave attenuata* Salm-Dyck | DBG 1983-0526-02-6 |
| *Agave bovicornuta* Gentry | *Agave bovicornuta* Gentry | DBG 2017-0677-01-4 |
| *Agave bovicornuta* Gentry | *Agave bovicornuta* Gentry | DBG 2019-0123-01-1 |
| *Agave bracteosa* S.Watson ex Englm. | *Agave bracteosa* S.Watson ex Englm. | DBG 1984-0108-01-1 |
| *Agave cerulata* subsp. *nelsonii* (Trel.) Gentry | *Agave cerulata* subsp. *nelsonii* (Trel.) Gentry | DBG 1939-0044-01-1 |
| *Agave deserti* Engelm. | *Agave deserti* Engelm. | DBG 1984-0119-21-13 |
| *Agave x glomeruliflora* (Engelm.) A.Berger | *Agave x glomeruliflora* (Engelm.) A.Berger | DBG 1963-7484-02-2 |
| *Agave murpheyi* Gibson | *Agave murpheyi* Gibson | DBG 2002-0337-01-1 |
| *Agave ocahui* Gentry | *Agave ocahui* Gentry | DBG 2004-0149-01-3 |
| *Agave palmeri* Engelm. | *Agave palmeri* Engelm. | DBG 1996-0062-02-6 |
| *Agave parryi* Engelm. | *Agave parryi* Engelm. | DBG 1990-0391-02-4 |
| *Agave scabra* Ortega | *Agave scabra* Ortega | DBG 1961-6898-01-2 |
| *Agave schotti* Engelm. | *Agave schotti* Engelm. | DBG 1985-0923-21-4 |

| | | |
|---|---|---|
| *Agave tequilana* F.A.C. Weber | *Agave tequilana* F.A.C. Weber | DBG 1978-0505-02-3 |
| *Alluaudia Dumosa* (Drake) Drake | *Alluaudia Dumosa* (Drake) Drake | DBG 1974-0228-01-1 |
| *Ariocarpus retusus* Scheidw. | *Ariocarpus retusus* Scheidw. | DBG 2018-0076-01-4 |
| *Astrophytum myriostigma* Lem. | *Astrophytum myriostigma* Lem. | DBG 1997-02443-01-3 |
| *Austrocylindropuntia subulata* (Muehlenpf.) Backeb. | *Austrocylindropuntia subulata* (Muehlenpf.) Backeb. | DBG 2018-0265-01-1 |
| *Beaucarnea recurvata* (K.Koch & Fintelm.) Lem. | *Beaucarnea recurvata* (K.Koch & Fintelm.) Lem. | DBG 1990-0232-01-1 |
| *Calymmanthium substerile* F.Ritter | *Calymmanthium substerile* F.Ritter | DBG 2015-0939-01-5 |
| *Carnegiea gigantea* (Engelm.) Britton & Rose | *Carnegiea gigantea* (Engelm.) Britton & Rose | DBG 1978-0542-01-19 |
| *Copiapoa rupestris* F.Ritter | *Copiapoa rupestris* F.Ritter | DBG 2014-2292-01-2 |
| *Cylindropuntia arbuscula* (Engelm.) F.M. Knuth | *Cylindropuntia arbuscula* (Engelm.) F.M. Knuth | DBG 2016-0433-01-1 |
| *Cylindropuntia imbricata* (Haw.) F.M. Knuth | *Cylindropuntia imbricata* (Haw.) F.M. Knuth | DBG 1967-8828-01-1 |
| *Cylindropuntia spinosior* (Engelm.) F.M. Knuth | *Cylindropuntia imbricata* subsp. *spinosior* (Engelm.) M.A. Baker, Cloud-H. & Majure | DBG No accession number |
| *Dasylirion wheeleri* S.Watson ex Rothr. | *Dasylirion wheeleri* S.Watson ex Rothr. | DBG 1977-0920-01-1 |
| *Didierea trolli* Capuron & Rauh | *Didierea trolli* Capuron & Rauh | DBG 1981-0024-02-2 |
| *Dracaena serrulata* Baker | *Dracaena serrulata* Baker | DBG 2009-0175-01-1 |
| *Echinocereus berlandieri* (Engelm.) Haage | *Echinocereus berlandieri* (Engelm.) Haage | DBG 1992-0429-01-1 |
| *Echinopsis aurea* Britton & Rose | *Echinopsis aurea* Britton & Rose | DBG 1950-2357-01-3 |
| *Echinopsis terscheckii* (J.Parm ex Pfeiff.) H.Friedrich & G.D.Rowley | *Leucostele terscheckii* (J.Parm ex Pfeiff.) Schlumpb. | DBG 1995-0042-10-22 |
| *Eriocyse subgibbosa* subsp. *nigrihorrida* (Backeb.) Katt. | *Eriocyse nigrihorrida* (Backeb.) P.C.Guerrero & Helmut Walter | DBG 2014-2006-01-3 |
| *Ferocactus latispinus* (Haw.) Britton & Rose | *Ferocactus latispinus* (Haw.) Britton & Rose | DBG 1986-0539-10-3 |
| *Ferocactus latispinus* (Haw.) Britton & Rose | *Ferocactus latispinus* (Haw.) Britton & Rose | DBG 2013-0447-01-1 |
| *Furcraea cahum* Trel. | *Fucraea hexapetala* (Jacq.) Urb | DBG 1987-0847-02-22 |
| *Furcraea macdougallii* Matuda | *Furcraea macdougallii* Matuda | DBG 2013-1074-01-3 |
| *Grusonia bradtiana* (J.M.Coult.) Britton & Rose | *Grusonia bradtiana* (J.M.Coult.) Britton & Rose | DBG 2014-0433-01-2 |
| *Gymnocalycium mihanovichii (Frič & Gürke) Britton & Rose* | *Gymnocalycium mihanovichii* (Frič & Gürke) Britton & Rose | DBG 2015-0269-01-1 |
| *Hesperaloe funifera* (K.Koch) Trel. | *Hesperaloe funifera* (K.Koch) Trel. | DBG 1976-0106-01-8 |

| | | |
|---|---|---|
| *Hesperaloe nocturna* (Gentry) | *Hesperaloe nocturna* (Gentry) | DBG 1991-0478-01-1 |
| *Hesperoyucca whipplei* (Torr.) Trel. | *Hesperoyucca whipplei* (Torr.) Trel. | DBG 2000-0044-10-8 |
| *Hylocereus lemairei* (Hook.) Britton & Rose | *Selenicereus monacanthus* (Lem.) D.R.Hunt | DBG 1999-0074-01-1 |
| *Lophocereus gatesii* M.E.Jones | *Lophocereus gatesii* M.E.Jones | DBG 1939-0269-02-1 |
| *Lophocereus schottii* (Engelm.) Britton & Rose | *Lophocereus schottii* (Engelm.) Britton & Rose | DBG 1979-0340-01-1 |
| *Manfreda undulata* (Klotzsch) Rose | *Agave undulata* Klotzsch | DBG 2002-0269-01-3 |
| *Manfreda virginica* (L.) Salisb. ex Rose | *Agave virginica* L. | DBG 1991-0455-01-1 |
| *Myrtillocactus geometrizans* (Mart. ex Pfeiff.) Console | *Myrtillocactus geometrizans* (Mart. ex Pfeiff.) Console | DBG 2018-0047-01-1 |
| *Nolina bigelovii* (Torr.) S.Watson | *Nolina bigelovii* (Torr.) S.Watson | DBG 2018-0626-01-3 |
| *Nolina matapensis* Wiggins | *Nolina matapensis* Wiggins | DBG 2018-0474-01-1 |
| *Opuntia bravoana* E.M.Baxter | *Opuntia bravoana* E.M.Baxter | DBG 2015-0303-01-1 |
| *Opuntia cochenillifera* (L.) Mill. | *Opuntia cochenillifera* (L.) Mill. | DBG 1997-0395-02-9 |
| *Opuntia gaumeri* (Britton & Rose) R.Puente & Majure | *Opuntia inaperta* (Schott ex Griffiths) | DBG 1997-0367-02-3 |
| *Opuntia robusta* H.L.Wendl. ex Pfeiff. | *Opuntia robusta* H.L.Wendl. ex Pfeiff. | DBG 1987-0849-21-3 |
| *Opuntia salmiana* J.Parm. ex Pfeiff. | *Salmonopuntia salmiana* (J.Parm. ex Pfeiff.) P.V.Heath | DBG 2001-0048-01-2 |
| *Peniocereus cuixmalensis* Sánchez-Mej. | *Acanthocereus cuixmalensis* (Sánchez-Mej.) Lodé | DBG 2004-0361-01-5 |
| *Pereskia sacharosa* (Griseb.) | *Rhodocactus sacharosa* (Griseb.) Backeb.[a] | DBG 1987-0471-21-10 |
| *Pereskiopsis porter* (Brandegee ex F.A.C.Weber) Britton & Rose | *Pereskiopsis porter* (Brandegee ex F.A.C.Weber) Britton & Rose | DBG 194-0537-01-2 |
| *Pterocactus tuberosus* (Pfeiff.) Britton & Rose | *Pterocactus tuberosus* (Pfeiff.) Britton & Rose | DBG 2013-1098-01-1 |
| *Quiabentia verticillate* (Vaupel) Borg | *Quiabentia verticillate* (Vaupel) Borg | DBG 1985-0461-01-1 |
| *Quiabentia verticillate* (Vaupel) Borg | *Quiabentia verticillate* (Vaupel) Borg | DBG 2013-1099-01-2 |
| *Sansevieria erythraeae* Mattei | *Dracaena erythraeae* (Mattei) Byng & Christenh. | DBG 2014-0193-02-2 |
| *Sansevieria Hyacinthoides* (L.) Druce | *Dracaena Hyacinthoides* (L.) Mabb. | DBG 1990-0419-02-1 |
| *Stenocereus alamosensis* (J.M.Coult.) A.C.Gibson & K.E.Horak | *Stenocereus alamosensis* (J.M.Coult.) A.C.Gibson & K.E.Horak | DBG 2016-0655-01-1 |
| *Stenocereus thurberi* (Engelm.) Buxb. | *Stenocereus thurberi* (Engelm.) Buxb. | DBG 1940-0425-01-1 |
| *Stetsonia coryne* (C.F.Först.) Britton & Rose | *Stetsonia coryne* (C.F.Först.) Britton & Rose | DBG 1980-0084-01-1 |
| *Tacinga lilae* (Trujillo & Marisela Ponce) Majure & R.Puente | *Tacinga lilae* (Trujillo & Marisela Ponce) Majure & R.Puente | DBG 1997-0369-21-1 |

| | | |
|---|---|---|
| *Tephrocactus articulates* (Pfeiff.) Backeb. | *Tephrocactus articulates* (Pfeiff.) Backeb. | DBG 1953-4443-01-1 |
| *Tunilla corrugate* (Salm-Dyck) D.R.Hunt & Iliff | *Airampoa corrugate* (Salm-Dyck) Doweld | DBG 2001-0006-01-1 |
| *Yucca x baccata* (Torr.) | *Yucca x baccata* (Torr.) | DBG 1997-0431-01-1 |
| *Yucca brevifolia* Engelm. | *Yucca brevifolia* Engelm. | DBG 1940-0478-01-1 |
| *Yucca brevifolia* Engelm. | *Yucca brevifolia* Engelm. | DBG 1976-0104-01-6 |
| *Yucca capensis* L.W.Lenz | *Yucca capensis* L.W.Lenz | DBG 2016-0804-01-3 |
| *Yucca constricta* Buckley | *Yucca constricta* Buckley | DBG 2014-0820-01-3 |
| *Yucca elata* (Engelm.) Engelm. | *Yucca elata* (Engelm.) Engelm. | DBG 1965-7784-01-1 |
| *Yucca glauca* Nutt. | *Yucca glauca* Nutt. | DBG 2009-0173-01-2 |
| *Yucca linearifolia* Clary | *Yucca linearifolia* Clary | DBG 2019-0251-01-2 |
| *Yucca pallida* McKelvey | *Yucca pallida* McKelvey | DBG 2014-0234-01-6 |
| *Yucca queretaroensis* Piña Luján | *Yucca queretaroensis* Piña Luján | DBG 2016-0006-01-1 |
| *Yucca rostrata* Engelm. ex Trel. | *Yucca rostrata* Engelm. ex Trel. | DBG 2017-0048-01-2 |
| *Yucca schidigera* Roezl ex Ortgies | *Yucca schidigera* Roezl ex Ortgies | DBG 2009-0202-01-1 |
| *Yucca x schottii* Engelm. | *Yucca x schottii* Engelm. | DBG 2007-0137-10-9 |
| *Yucca torreyi* Shafer | *Yucca treculiana* Carrière | DBG 2002-0163-10-2 |
| *Alluaudia procera* (Drake) Drake | *Alluaudia procera* (Drake) Drake | No accession number |
| *Anacampseros lanceolata* (Haw.) Sweet | *Anacampseros lanceolata* (Haw.) Sweet | AJM 1635 |
| *Anacampseros rufescens* (Haw.) Sweet | *Anacampseros rufescens* (Haw.) Sweet | AJM 1641 |
| *Anredera baselloides* (Knuth) Baill. | *Anredera baselloides* (Knuth) Baill. | AJM 1583 |
| *Calandrinia brevipedata* F.Muell. | *Parakeelya brevipedata* (F.Muell.) Hershk.[b] | LPH 50A |
| *Calandrinia flava* Obbens | *Parakeelya flava* (Obbens) Hershk.[b] | JAH 6 |
| *Calandrinia kalanniensis* Obbens | *Parakeelya kalanennsis* (Obbens) Hershk.[b] | LPH 143 |
| *Calandrinia liniflora* Finzl | *Parakeelya linliflora* (Finzl) Hershk.[b] | LPH 49A |
| *Calandrinia pentavalvis* Obbens | *Parakeelya pentavalvis* (Obbens) Hershk.[b] | LPH 147 |
| *Calandrinia pleiopetala* F.Muell. | *Parakeelya pleiopetala* (F.Muell) Hershk.[b] | JAH 3 |
| *Calandrinia pumila* (Benth.) F.Muell. | *Parakeelya pumila* (F.Muell) Hershk.[b] | LPH 173 |
| *Calandrinia quadrivalvis* F.Muell | *Parakeelya quadrivalvus* (F.Muell) Hershk.[b] | LPH 156 |
| *Calandrinia schistorhiza* Morrison | *Parakeelya schistorhiza* (Morrison) Hershk.[b] | LPH 149 |
| *Calandrinia spergularina* F.Muell. | *Parakeelya spergularina* (F.Muell) Hershk.[b] | LPH 133 |
| *Calandrinia tumida* Syeda | *Parakeelya tumida* (Syeda) Hershk.[b] | JAH 138 |

[a]Because of phylogenetic uncertainty, we maintain the use of *"Pereskia*", rather than *Leuenbergeria* and *Rhodocactus*, but recognize that *Pereskia* is non-monophyletic.
[b]We prefer the use of *Parakeelya* to for the Australian members of the paraphyletic genus *Calandrinia*, following Theiele *et al.* (2018).

**Table S3** Results of D'Angostino and Pearson's test for normality and Bartlett's test for homoscedasticity of raw and $log_{10}$-transformed data.

| Test | Data | Mesophyll cell area (μm²) | Leaf thickness (μm) | Intercellular airspace (%) | Leaf dry matter content (g/g) | Specific leaf area (mm²/mg dry mass) |
|---|---|---|---|---|---|---|
| Normality | raw | $p = 5.64 \times 10^{-52}$ | $p = 0$ | $p = 9.05 \times 10^{-7}$ | $p = 5.65 \times 10^{-27}$ | $p = 0$ |
| Normality | $log_{10}$ | $p = 2.93 \times 10^{-5}$ | $p = 3.55 \times 10^{-65}$ | $p = 5.95 \times 10^{-18}$ | $p = 2.95 \times 10^{-25}$ | $p = 2.07 \times 10^{-20}$ |
| Homoscedasticity | raw | $p = 4.14 \times 10^{-64}$ | $p = 1.01 \times 10^{-245}$ | $p = 1.75 \times 10^{-6}$ | $p = 1.75 \times 10^{-13}$ | $p = 2.11 \times 10^{-34}$ |
| Homoscedasticity | $log_{10}$ | $p = 0.1589$ | $p = 0.7650$ | $p = 0.5634$ | $p = 0.7417$ | $p = 0.0003$ |

**Table S4** Node calibrations used from Arakaki *et al.* (2011).

| Node | Age (Ma) |
|---|---|
| Opuntioideae | 15.2 |
| Portulacaceae | 17.6 |
| Didieroideae | 18.0 |
| Cactoideae | 19.7 |
| Core Cactaceae | 25.0 |
| Cactaceae | 28.0 |
| Talinaceae | 29.9 |
| Anacampserotaceae + Cactaceae + Portulacaceae (ACP) | 36.9 |
| ACP + Talinaceae (ACPT) | 39.4 |
| Montiaceae | 39.9 |
| ACPT + Basellaceae + Didiereaceae + Montiaceae + Talinaceae (Portulacineae) | 44.9 |
| Molluginaceae | 46.2 |
| Molluginaceae + Portulacineae (Portullugo) | 53.0 |

**Table S5** Results of Kruskal–Wallis tests for group difference between CAM phenotypes.

| Feature | Test result |
|---|---|
| Intercellular airspace (IAS) | $H = 44.1$ ($p = 2.66 \times 10^{-10}$) |
| Leaf dry matter content (LDMC) | $H = 151.7$ ($p = 1.13 \times 10^{-33}$) |
| Leaf thickness (LT) | $H = 715.0$ ($p = 5.52 \times 10^{-156}$) |
| Mesophyll cell size (MA) | $H = 208.9$ ($p = 4.40 \times 10^{-46}$) |
| Specific leaf area (SLA) | $H = 57.3$ ($p = 3.69 \times 10^{-13}$) |

**Table S6** Relative feature importance of multiclass models.

| Model | LT | MA | LDMC | SLA | IAS |
|---|---|---|---|---|---|

| | | | | | |
|---|---|---|---|---|---|
| DART | 33.725% | 28.105% | 13.203% | 10.196% | 14.771% |
| DART+ROS | 31.404% | 23.794% | 16.613% | 15.434% | 12.755% |
| DART+RUS | 32.839% | 24.675% | 12.245% | 20.037% | 10.204% |
| DART+Iter. | 19.218% | 20.847% | 18.893% | 14.115% | 26.927% |
| DART+Knn | 20.813% | 22.727% | 20.813% | 14.593% | 21.053% |
| DART+Med. | 34.113% | 33.287% | 10.867% | 8.803% | 12.930% |
| DART+MDS1 | 35.188% | 27.296% | 11.902% | 10.737% | 14.877% |
| DART+MDS2 | 33.725% | 28.105% | 13.203% | 10.196% | 14.771% |
| DART+MDS5 | 33.725% | 28.105% | 13.203% | 10.196% | 14.771% |
| DART+MDS10 | 33.725% | 28.105% | 13.203% | 10.196% | 14.771% |
| gbtree | 32.238% | 26.043% | 14.918% | 12.389% | 14.412% |
| gbtree+ROS | 32.576% | 23.052% | 17.208% | 14.935% | 12.229% |
| gbtree+RUS | 33.152% | 24.094% | 12.862% | 17.935% | 11.957% |
| gbtree+Iter. | 20.444% | 23.222% | 17.556% | 14.444% | 24.333% |
| gbtree+Knn | 22.432% | 23.849% | 20.071% | 13.813% | 19.835% |
| gbtree+Med. | 32.510% | 31.139% | 13.580% | 11.248% | 11.523% |
| gbtree+MDS1 | 32.826% | 26.869% | 14.068% | 11.914% | 14.322% |
| gbtree+MDS2 | 32.238% | 26.043% | 14.918% | 12.389% | 14.412% |
| gbtree+MDS5 | 32.238% | 26.043% | 14.918% | 12.389% | 14.412% |
| gbtree+MDS10 | 32.238% | 26.043% | 14.918% | 12.389% | 14.412% |
| Mean ± SD | 30.6 ± 5% | 26.1 ± 2.9% | 15 ± 2.7% | 12.9 ± 2.7% | 15.5 ± 4.2% |

**Table S7** Incorrect predictions of the best performing multiclass model.

Incorrect predictions of the best performing multiclass model (gbtree+ROS) from one representative sample of testing data.

| Major Lineage | Taxon | Pathway | True label | Predicted label |
|---|---|---|---|---|
| Acanthaceae | *Blepharis ciliaris* | C4 | nonCAM | mCAM |
| Amaranthaceae | *Suaeda vera* | C3 | nonCAM | pCAM |
| Asteraceae | *Baccharis articulata* | C3 | nonCAM | pCAM |
| Asteraceae | *Noticastrum marginatum* | C3 | nonCAM | mCAM |
| Bromeliaceae | *Catopsis paniculata* | C3 | nonCAM | pCAM |
| Bromeliaceae | *Guzmania claviformis* | C3 | nonCAM | pCAM |
| Bromeliaceae | *Guzmania sanguinea* | C3 | nonCAM | pCAM |
| Bromeliaceae | *Racinaea aeris-incola* | C3 | nonCAM | mCAM |
| Bromeliaceae | *Vriesea fenestralis* | C3 | nonCAM | mCAM |
| Caryophyllaceae | *Arenaria lychnidea* | C3 | nonCAM | mCAM |
| Cyperaceae | *Carex rostrata* | C3 | nonCAM | mCAM |
| Ericaceae | *Macleania rupestris* | C3 | nonCAM | mCAM |
| Euphorbiaceae | *Macaranga gigantea* | C3 | nonCAM | mCAM |

| | | | | |
|---|---|---|---|---|
| Fabaceae | *Acacia havilandiorum* | C3 | nonCAM | pCAM |
| Fabaceae | *Senna aphylla* | C3 | nonCAM | pCAM |
| Fabaceae | *Ulex europaeus* | C3 | nonCAM | mCAM |
| Fagaceae | *Lithocarpus hancei* | C3 | nonCAM | mCAM |
| Lamiaceae | *Salvia verticillata* | C3 | nonCAM | mCAM |
| Molluginaceae | *Hypertelis salsoloides* | C3 | nonCAM | pCAM |
| Molluginaceae | *Mollugo verticillata* | C3-C4 | nonCAM | mCAM |
| Montiaceae | *Claytonia megarhiza* | C3 | nonCAM | mCAM |
| Montiaceae | *Montia parvifolia* | C3 | nonCAM | mCAM |
| Montiaceae | *Montiopsis andicola* | C3 | nonCAM | mCAM |
| Moraceae | *Antiaris africana* | C3 | nonCAM | mCAM |
| Nolinoideae | *Nolina matapensis* | C3 | nonCAM | mCAM |
| Orchidaceae | *Arundina graminifolia* | C3 | nonCAM | mCAM |
| Orchidaceae | *Calanthe sp. 1* | C3 | nonCAM | mCAM |
| Orchidaceae | *Ceratostylis sp. 13* | C3 | nonCAM | mCAM |
| Orchidaceae | *Ceratostylis sp. 3* | C3 | nonCAM | pCAM |
| Orchidaceae | *Draconia tuerckheimii* | C3 | nonCAM | mCAM |
| Orchidaceae | *Galeottia grandiflora* | C3 | nonCAM | mCAM |
| Orchidaceae | *Masdevallia zahlbruckneri* | C3 | nonCAM | mCAM |
| Orchidaceae | *Maxillaria tenuifolia* | C3 | nonCAM | mCAM |
| Orchidaceae | *Oncidium parviflorum* | C3 | nonCAM | mCAM |
| Orchidaceae | *Orchis mascula* | C3 | nonCAM | mCAM |
| Orchidaceae | *Pleurothallis crocodiliceps* | C3 | nonCAM | pCAM |
| Orchidaceae | *Pleurothallis sp.* | C3 | nonCAM | pCAM |
| Orchidaceae | *Sobralia chrysostoma* | C3 | nonCAM | mCAM |
| Orchidaceae | *Specklinia fulgens* | C3 | nonCAM | pCAM |
| Orchidaceae | *Warczewiczella lipscombiae* | C3 | nonCAM | mCAM |
| Orchidaceae | *Zygopetalum intermedii* | C3 | nonCAM | mCAM |
| Plantaginaceae | *Kickxia spuria* | C3 | nonCAM | mCAM |
| Plantaginaceae | *Plantago australis* | C3 | nonCAM | mCAM |
| Plantaginaceae | *Veronica arvensis* | C3 | nonCAM | mCAM |
| Poaceae | *Brachypodium retusum* | C3 | nonCAM | mCAM |
| Polygonaceae | *Polygonum persicaria* | C3 | nonCAM | mCAM |
| Polypodiaceae | *Lepisorus spicatus* | C3 | nonCAM | pCAM |
| Polypodiaceae | *Lepisorus validinervis* | C3 | nonCAM | mCAM |
| Polypodiaceae | *Selliguea plantaginea* | C3 | nonCAM | mCAM |
| Primulaceae | *Hottonia palustris* | C3 | nonCAM | mCAM |
| Santalaceae | *Exocarpos aphyllus* | C3 | nonCAM | pCAM |
| Saxifragaceae | *Saxifraga hirsuta* | C3 | nonCAM | mCAM |

| | | | | |
|---|---|---|---|---|
| Solanaceae | *Nicotiana glauca* | C3 | nonCAM | mCAM |
| Theaceae | *Camellia japonica* | C3 | nonCAM | mCAM |
| Theaceae | *Gordonia axillaris* | C3 | nonCAM | mCAM |
| Viburnaceae | *Sambucus nigra* | C3 | nonCAM | pCAM |
| Bromeliaceae | *Tillandsia complanata* | C3+CAM | mCAM | nonCAM |
| Bromeliaceae | *Werauhia sanguinolenta* | C3+CAM | mCAM | nonCAM |
| Cactaceae | *Pereskia marcanoi* | C3+CAM | mCAM | nonCAM |
| Cactaceae | *Pereskia weberiana* | C3+CAM | mCAM | nonCAM |
| Crassulaceae | *Sedum acre* | C3+CAM | mCAM | pCAM |
| Crassulaceae | *Sedum album* | C3+CAM | mCAM | nonCAM |
| Didiereaceae | *Ceraria namaquensis* | C3+CAM | mCAM | nonCAM |
| Gesneriaceae | *Codonanthe uleana* | C3+CAM | mCAM | nonCAM |
| Montiaceae | *Cistanthe picta* | C3+CAM | mCAM | pCAM |
| Montiaceae | *Cistanthe salsoloides* | C3+CAM | mCAM | pCAM |
| Orchidaceae | *Echinosepala lappiformis* | C3+CAM | mCAM | pCAM |
| Orchidaceae | *Epidendrum rousseauae* | C3+CAM | mCAM | pCAM |
| Orchidaceae | *Lockhartia micrantha* | C3+CAM | mCAM | nonCAM |
| Orchidaceae | *Oncidium dichromaticum* | C3+CAM | mCAM | nonCAM |
| Portulacaceae | *Portulaca bicolor* | C4+CAM | mCAM | pCAM |
| Agavoideae | *Agave cerulata* | Primary CAM | pCAM | mCAM |
| Apocynaceae | *Dischidia chinensis* | Primary CAM | pCAM | nonCAM |
| Bromeliaceae | *Aechmea capixabae* | Primary CAM | pCAM | mCAM |
| Bromeliaceae | *Lymania globosa* | Primary CAM | pCAM | mCAM |
| Crassulaceae | *Crassula deceptor* | Primary CAM | pCAM | nonCAM |
| Orchidaceae | *Dendrobium lineale* | Primary CAM | pCAM | mCAM |
| Orchidaceae | *Encyclia stellata* | Primary CAM | pCAM | nonCAM |
| Orchidaceae | *Notylia pentachne* | Primary CAM | pCAM | nonCAM |
| Orchidaceae | *Vanilla fragrans* | Primary CAM | pCAM | mCAM |
| Orchidaceae | *Vanilla inodora* | Primary CAM | pCAM | mCAM |

**Table S8** Incorrect predictions of the best performing binary model.

Incorrect predictions of the best performing binary model (gbtree+hinge+ROS) from one representative sample of testing data.

| Major Lineage | Taxon | Pathway | True label | Predicted label |
|---|---|---|---|---|
| Agavoideae | *Yucca constricta* | C3 | nonCAM | CAM |
| Bromeliaceae | *Guzmania sanguinea* | C3 | nonCAM | CAM |
| Bromeliaceae | *Lutheria glutinosa* | C3 | nonCAM | CAM |

| | | | | |
|---|---|---|---|---|
| Bromeliaceae | *Puya mirabilis* | C3 | nonCAM | CAM |
| Bromeliaceae | *Vriesea gigantea* | C3 | nonCAM | CAM |
| Bromeliaceae | *Vriesea pleiosticha* | C3 | nonCAM | CAM |
| Clusiaceae | *Clusia multiflora* | C3 | nonCAM | CAM |
| Clusiaceae | *Tomovita lanceolata* | C3 | nonCAM | CAM |
| Fabaceae | *Gompholobium glabratum* | C3 | nonCAM | CAM |
| Fabaceae | *Ulex europaeus* | C3 | nonCAM | CAM |
| Malvaceae | *Lueheopsis rugosa* | C3 | nonCAM | CAM |
| Molluginaceae | *Pharnaceum microphyllum* | C3 | nonCAM | CAM |
| Montiaceae | *Claytonia megarhiza* | C3 | nonCAM | CAM |
| Nolinoideae | *Nolina bigelovii* | C3 | nonCAM | CAM |
| Nyctaginaceae | *Mirabilis nyctaginea* | C3 | nonCAM | CAM |
| Orchidaceae | *Acianthera johnsonii* | C3 | nonCAM | CAM |
| Orchidaceae | *Ceratostylis sp. 13* | C3 | nonCAM | CAM |
| Orchidaceae | *Ceratostylis sp. 4* | C3 | nonCAM | CAM |
| Orchidaceae | *Cyrtochiloides ochmatochila* | C3 | nonCAM | CAM |
| Orchidaceae | *Oncidium cheirophorum* | C3 | nonCAM | CAM |
| Orchidaceae | *Oncidium leucochilum* | C3 | nonCAM | CAM |
| Orchidaceae | *Pleurothallis sp.* | C3 | nonCAM | CAM |
| Orchidaceae | *Scaphyglottis behrii* | C3 | nonCAM | CAM |
| Orchidaceae | *Sobralia macrophylla* | C3 | nonCAM | CAM |
| Orchidaceae | *Stanhopea oculata* | C3 | nonCAM | CAM |
| Orchidaceae | *Zygopetalum intermedii* | C3 | nonCAM | CAM |
| Primulaceae | *Primula veris* | C3 | nonCAM | CAM |
| Sapotaceae | *Pouteria grandis* | C3 | nonCAM | CAM |
| Solanaceae | *Physalis pumila* | C3 | nonCAM | CAM |
| Theaceae | *Gordonia axillaris* | C3 | nonCAM | CAM |
| Typhaceae | *Typha latifolia* | C3 | nonCAM | CAM |
| Cactaceae | *Pereskia weberiana* | C3+CAM | CAM | nonCAM |
| Crassulaceae | *Crassula helmsii* | C3+CAM | CAM | nonCAM |
| Gesneriaceae | *Codonanthe uleana* | C3+CAM | CAM | nonCAM |
| Montiaceae | *Parakeelya schistorhiza* | C3+CAM | CAM | nonCAM |
| Orchidaceae | *Coryanthes hunteriana* | C3+CAM | CAM | nonCAM |
| Orchidaceae | *Peristeria sp.* | C3+CAM | CAM | nonCAM |
| Orchidaceae | *Saccolabium sp. 1* | Primary CAM | CAM | nonCAM |
| Orchidaceae | *Scaphyglottis imbricata* | C3+CAM | CAM | nonCAM |

**Table S9** Phylogenetic signal in anatomical features of the Portullugo.

| Feature | Blomberg's $K$ | Pagel's $\lambda$ |
|---|---|---|
| Intercellular airspace (IAS) | 0.58 ($p = 0.026$) | 0.56 ($p = 0.005$) |
| Leaf thickness (LT) | 1.09 ($p < 0.001$) | 0.91 ($p < 0.001$) |
| Mesophyll cell size (MA) | 1.15 ($p < 0.001$) | 0.89 ($p < 0.001$) |

**Table S10** Results of phylogenetic least squares (PGLS) regressions.

Underlying models of trait evolution are given in parentheses; "*" indicate slopes significantly different from 0 ($p < 0.05$); bolded models have significant slopes and were selected as best fit using AIC values; all trait values are $log_{10}$-transformed; MA, mesophyll cell area; LT, leaf thickness; IAS, intercellular airspace.

| PGLS model | Intercept (± SE) | Slope (± SE) | AIC | DF |
|---|---|---|---|---|
| **IAS ~ MA (BM)** | -1.82 ± 0.37 | 0.29 ± 0.10* | 8.058725 | 49 |
| IAS ~ MA (OU) | -1.37 ± 0.35 | 0.15 ± 0.10 | 3.815509 | 49 |
| **LT ~ MA (BM)** | 1.52 ± 0.38 | 0.39 ± 0.11* | 8.357876 | 39 |
| **LT ~ MA (OU)** | **1.13 ± 0.29** | **0.51 ± 0.08*** | **-5.06187** | **39** |
| LT ~ IAS (BM) | 2.91 ± 0.22 | 0.10 ± 0.20 | 22.5635 | **35** |
| LT ~ IAS (OU) | 3.02 ± 0.20 | 0.21 ± 0.21 | 20.69011 | 35 |
| MA ~ CAM phenotype (BM) | 3.19 ± 0.28 | 0.08 ± 0.11 | 76.52515 | 68 |
| **MA ~ CAM phenotype (OU)** | **3.13 ± 0.13** | **0.25 ± 0.08*** | **44.35982** | **68** |
| LT ~ CAM phenotype (BM) | 2.66 ± 0.15 | 0.26 ± 0.10* | 8.6479 | 40 |
| **LT ~ CAM phenotype (OU)** | **2.66 ± 0.12** | **0.30 ± 0.09*** | **5.771516** | **40** |
| IAS ~ CAM phenotype (BM) | -0.62 ± 0.17 | -0.22 ± 0.08 | 9.412173 | 50 |
| **IAS ~ CAM phenotype (OU)** | **-0.65 ± 0.07** | **-0.20 ± 0.05*** | **-4.89543** | **50** |

**Methods S1** Summary of MiniContourFinder image segmentation algorithm

MiniContourFinder is a lightweight image segmentation tool built in Python v3 with OpenCV v4.5.2 (Bradski, 2000). MiniContourFinder was designed to allow users with minimal experience on the command line or with image processing to generate accurate and reproducible contours within minutes. MiniContourFinder can be run entirely through the command line or via a graphical user interface (GUI) with simple, adjustable parameters on sliders to tune the contours in real time. As a single parameter set may not be appropriate over the entirety of an image, the GUI implementation further allows the user to selectively save individual contours as they alter segmentation parameters. Contours, contour metadata (i.e., collection parameters), and a variety of shape metrics are exported in CSV and JSON format for readability as data frames in popular bioinformatics languages such as Python and R.

Segmentation in MiniContourFinder is accomplished through a combination of thresholding, gradient, and morphological operations (Fig. S1). The effects of these operations are governed by the kernel size over which they are applied, with smaller kernels placing higher weights on nearby pixels, and vice versa for larger kernels. The image (Fig. S1a) is first denoised using non-local means (Buades *et al.*, 2011), converted to grayscale, and undergoes adaptive histogram normalization to increase contrast (Fig. S1b). Then, the image is adaptively blurred using a low pass filter to augment the contiguity of boundaries (Fig. S1c) and an adaptive Gaussian threshold is applied to again increase the contrast of lines (Fig. S1d). A Laplacian operator acts as a high pass filter to sharpen the image (Fig. S1e), and the image is dilated to expand boundaries (Fig. S1f). The gradient is taken to remove space within hollow contours (Fig. S1g), and the result is binarized (Fig. S1h). Finally, the background (non-contour) is cleaned through morphological opening (erosion followed by dilation) (Fig. S1I) and closing (dilation followed by erosion) (Fig. S1j). To remove partial objects that extend beyond the image boundary, the image is flooded from the outside (Fig. S1k). Contours are then detected (Fig. S1l), with only the outermost contour returned in the case that contours are detected within one another.

MiniContourFinder can also detect and read scale bars within images. Scale bar detection is done through Canny edge detection (Canny, 1986) and a Hough line transform (Ballard, 1981). Scale bar units are then read using 'image_to_string' from pytesseract v0.3.0, a wrapper for the Tesseract optical character recognition (OCR) engine (Smith, 2007). Scale bar recognition is most successful when scale bars are placed away from the focal parts of the image and when images have few straight lines that may interfere with detection. If automated scale bar detection is unsuccessful, users can manually draw a scale bar to convert pixel-based measurements. MiniContourFinder can also approximate contours using convex hulls or approximate polygons (which allow for concave polygons) that can be useful for tasks where precise segmentations are not needed (e.g., counting) or when the desired segments are more regular.
